# Dynamic evolution in the key honey bee pathogen Deformed wing virus

**DOI:** 10.1101/653543

**Authors:** Eugene V. Ryabov, Anna K. Childers, Dawn Lopez, Kyle Grubbs, Francisco Posada-Florez, Daniel Weaver, William Girten, Dennis vanEngelsdorp, Yanping Chen, Jay D. Evans

## Abstract

The impacts of invertebrate RNA virus population dynamics on virulence and infection outcomes are poorly understood. Deformed wing virus (DWV), the main viral pathogen of honey bees, negatively impacts bee health which can lead to colony death. Despite previous reports on the reduction of DWV diversity following the arrival of the parasitic mite *Varroa destructor*, the key DWV vector, we found high genetic diversity of DWV in infested United States (US) honey bee colonies. Phylogenetic analysis showed that the divergent US DWV genotypes are of monophyletic origin, which were likely generated as a result of diversification after a genetic bottleneck. To investigate the population dynamics of this divergent DWV, we designed a series of novel infectious cDNA clones corresponding to co-existing DWV genotypes, thereby devising a reverse genetic system for an invertebrate RNA virus quasispecies. Equal replication rates were observed for all clone-derived DWV variants in single infections. Surprisingly, individual clones replicated to the same high levels as their mixtures and even the parental highly diverse natural DWV population, suggesting that complementation between genotypes was not required to replicate to high levels. Mixed clone-derived infections showed a lack of strong competitive exclusion, suggesting that the DWV genotypes were adapted to co-exist. Mutational and recombination events were observed across clone progeny providing new insights into the forces that drive and constrain virus diversification. Accordingly, herein we propose a new model of *Varroa*-induced DWV dynamics whereby an initial selective sweep is followed by virus diversification fueled by negative frequency-dependent selection for new genotypes. This selection likely reflects the ability of rare lineages to evade host defenses, specifically antiviral RNA interference (RNAi). In support of this, we show that RNAi induced against one DWV strain is less effective against an alternate strain from the same population.

**Author Summary:** Virulence of Deformed wing virus (DWV), a major pathogen of honey bees, showed a sharp and significant increase following the introduction of its vector, the mite *Varroa destructor*. *Varroa* vectoring resulted in genetic changes of DWV, including reduction of DWV diversity to nearly clonal levels in the UK and Hawaii. Contrary to the previous reports, we discovered that virulent DWV populations circulating across the *Varroa*-infested United States included many divergent genotypes generated following a strong bottleneck event. We designed a series of the full-length infectious cDNA clones that captured the diversity of a typical virulent DWV population from a declining *Varroa*-infested colony, effectively establishing first reverse genetic system for an invertebrate RNA virus quasispecies, in order to investigate interactions between the virus genotypes. We demonstrated that individual co-existing DWV genotypes and diverse natural DWV populations replicated equally well indicating that complementation between isolates was not required to enable DWV replication to high levels. Also, no obvious competitive exclusion was detected between genotypes in mixed infections suggesting DWV genotypes are adapted to co-exist to maintain overall population diversity. We suggest that introduction of *Varroa* resulted in an initial selective sweep of DWV diversity which was followed by DWV diversification driven by selection for new genotypes capable of evading host defenses, specifically antiviral RNA interference.

## Introduction

Population fluctuations affect virulence of invertebrate RNA viruses including the positve-strand RNA *Deformed wing virus* (DWV; Fig. 1A), a major pathogen of honey bees. DWV includes three master variants, the most widespread being DWV-A, followed by *Varroa destructor virus 1* (VDV1 or DWV-B), and infrequently, DWV-C [1–4]. Prevalance of DWV, tightly linked with its invasive vector the ectoparasitic mite *Varroa destructor* [5], increases honey bee colony mortality [6, 7], thereby threatening food security worldwide by affecting crop pollination [8]. Prior to *Varroa* invasion, DWV infections had low-titers and were non-symptomatic [9]. Indeed a phylogeographic study confirmed that the DWV pandemic and increase in virus virulence coincided with the global spread of the *Varroa* mite [10]. This suggests that DWV populations in *Varroa*-free and *Varroa-*infested colonies are genetically distinct, and that mite vectoring drives evolutionary changes in DWV. However, it is not clear which genetic changes make *Varroa*-transmitted DWV more virulent. Previous reports suggested that the VDV1-DWV recombinants detected in the UK and France [11–13] were associated with mite transmission, although similar recombinants were also present in UK *Varroa*-free honey bees [12]. The most striking difference found in DWV populations associated with mite transmission was a greatly reduced genetic diversity. This was observed in Hawaii, where a sudden drop of DWV diversity occurred following mite invasion [9]. Additionally in the UK, nearly clonal diversity of DWV-like viruses was reported in individual symptomatic *Varroa*-infested honey bees, contrasting with high viral variability in asymptomatic bees with low DWV levels [12]. Together these reports suggested that low genetic diversity could be a universal feature of the mite-transmitted DWV. Surprisingly, our analysis of the *Varroa*-associated DWV population currently circulating in the mainland United States (US) indicated high genetic diversity of DWV. To investigate the evolutionary dynamics of DWV, we designed a reverse-genetic system for a virulent DWV population by cloning variants co-existing in a typical US DWV-A quasispecies [14], the first of this kind for an RNA invertebrate virus. Using this system we demonstrated that replication levels of individual US DWV genotypes were equivalent to divergent wild-type DWV, without strong competitive exclusion in mixed infections. Recombination events between DWV isolates were widespread, contributing to virus diversification. We propose a model of DWV dynamics, potentially consistent with punctuated evolution [15], whereby introduction of *Varroa*-selected genotypes causes a selective sweep after which diversification *via* negative frequency-dependent selection results in high genetic heterogeneity, potentially benefiting the virus’s ability to escape genotype-specific antiviral defenses.

**Fig. 1.**
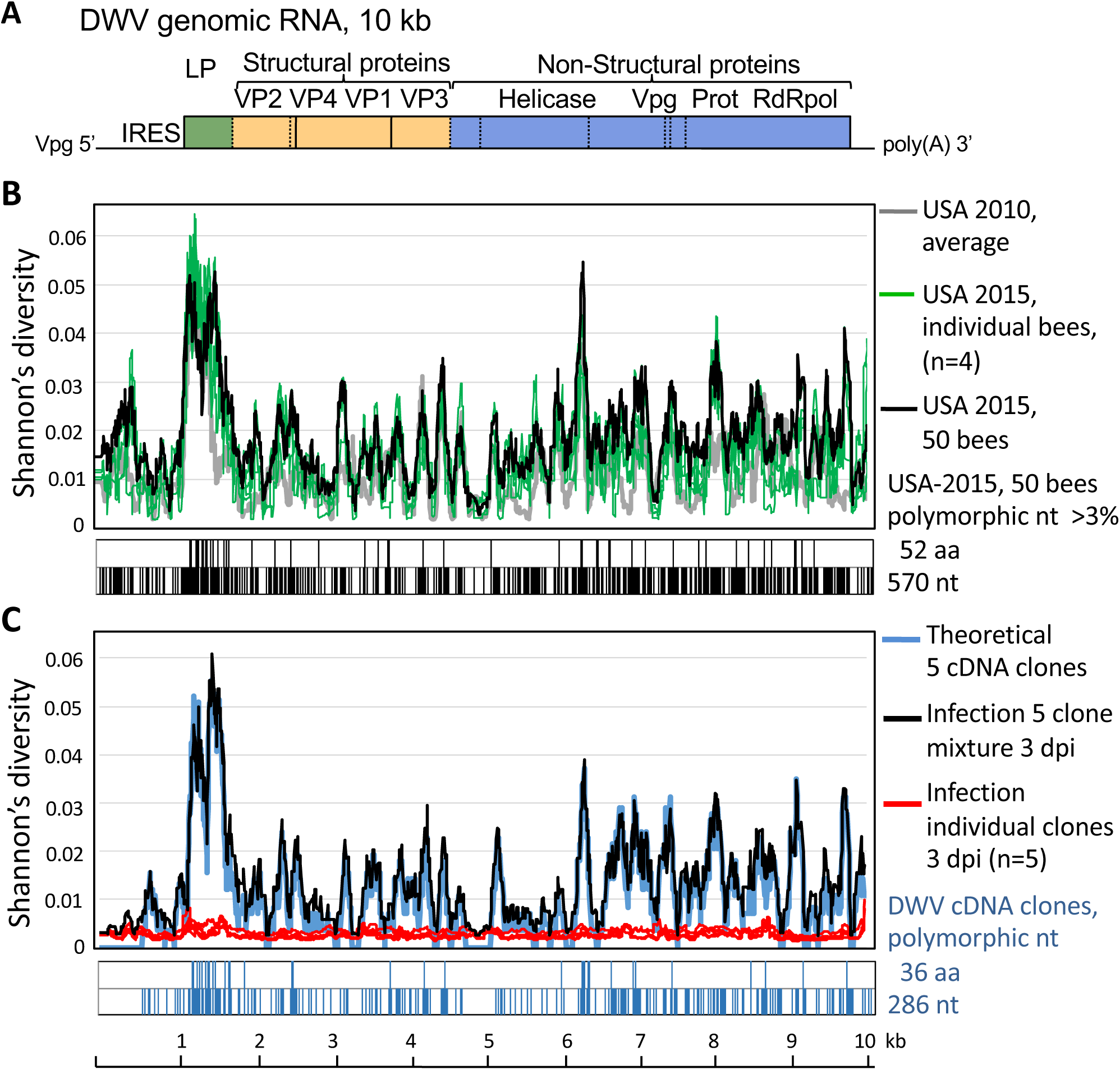
Polymorphisms in natural and clone-derived DWV. (A) Reference DWV of DWV genomic RNA. (B,C) Shannon’s diversity profiles for DWV (averaged for 100 nt) for (B) 2010 bees (grey line), 2015 individual honey bees (green lines) and a colony sample (black line); (C) individual bees infected with cDNA clone-derived DWV isolates, red lines – single infections, black line – mixed infection of five clones, blue line - theoretical profile for five clone mixture. Distributions of polymorphic nucleotides showing an alternate allele exceeding 3% in frequency and the resulting amino acid changes are shown below the diversity graphs.

## Results

### High genetic diversity of DWV in US *Varroa*-infested honey bee colonies reflects post-bottleneck expansion

Honey bee viromes comprehensively characterized by next generation sequencing (NGS) (S1 Table) had surprisingly high DWV genetic diversity in US *Varroa*-infested colonies compared to prior population surveys of the *Varroa*-associated DWV carried out in the United Kingdom (UK) in 2013 [12] (Fig. 2, S2 Table). In particular, the levels of DWV genetic diversity in individual US honey bees with high DWV levels (Fig, 2A, Group 4) were significantly higher than in overtly infected UK bees with high DWV levels [12] (Fig. 2A, Group 2) contradicting the previously reported nearly clonal nature of DWV in *Varroa*-infested colonies in the Hawaiian islands [9] and UK [12]. In fact, the US honey bees with both high and low DWV levels (Fig. 2A, Groups 3 and 4) showed virus diversity levels similar to those observed in covertly-infected UK bees with low DWV-levels (Fig. 2A, Groups 1). Significantly higher diversity level in the US honey bees compared to the UK bees was also observed at the colony level when the pools of 50 bees were analyzed by NGS. While UK colony-level DWV population was nearly clonal (Fig. 2A,B; “Colony UK 2013”, black arrows), the diversity of DWV in the US colony (Fig. 2A,B; “Colony USA 2015”, green arrow), was even higher than in the UK bees with low DWV levels suggesting that DWV populations in the US have more genetic variants.

**Fig. 2.**
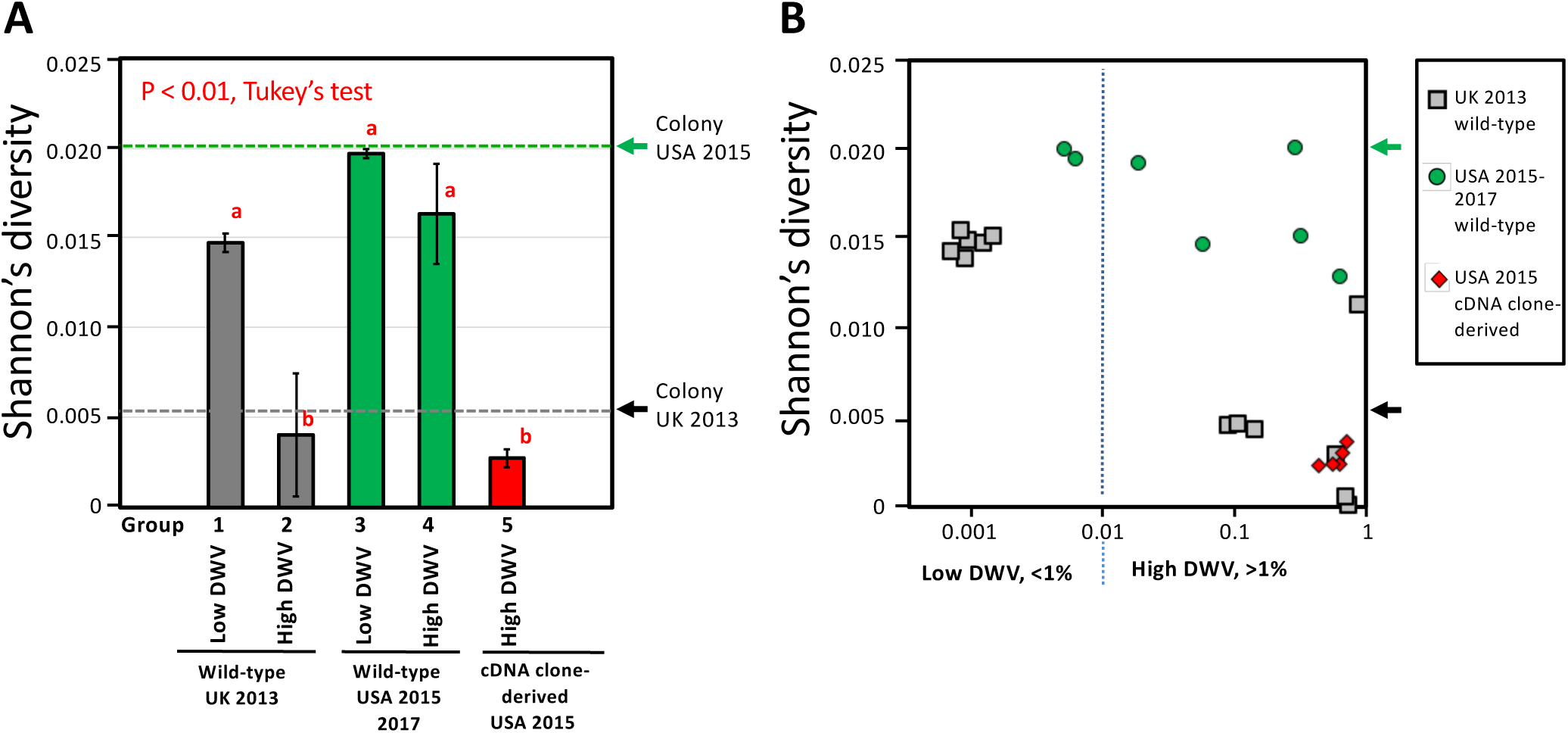
The connection between virus levels and in the US and the UK honey bees. (A) The columns show the average Shannon’s diversity index for the Non-Structural (NS) region of DWV genomic RNA for individual honey bees from the UK (UK 2013) and the USA (USA 2015) with low DWV or high DWV levels (less than 1 % or more than 1% of DWV reads in NGS library, correspondingly), and honey bees infected with the clone-derived US DWV variants. Error bars indicate standard deviation. The groups with significantly different (P<0.01) Shannon’s diversity index values are indicated above the bars with different letters. (B) Average Shannon’s diversity index for the NS region, plotted against the proportion of DWV in NGS library. (A, B) Arrows at the right side of the graphs indicate Shannon’s diversity index values of the colony-level DWV populations in the UK and the USA were calculated for the NGS libraries for virus preparations from 50 bees. The DWV NS region corresponds to the positions 5500-9847 of the US DWV isolate DWV-304, GenBank accession number MG831200.

Indeed, analysis of the divergent position distribution in a virulent DWV population from a US colony that collapsed three months after sampling, showed that it had 5.6% polymorphic nucleotides across the viral genome, where the proportion of the alternate nucleotide exceeded 3%, generating 1.8% amino acid substitutions (Fig. 1B, “USA 2015, 50 bees”; S1 Table, V99). NGS analysis also showed that this collapsed honey bee colony included exclusively DWV genotypes of type A (DWV-A), most closely related to US isolates DWV-PA [1] and DWV-Ame711 [16] and did not harbor VDV1 type, which has spread rapidly in the US in the last decade [17]. DWV polymorphism levels and distribution of diversity in individual honey bee pupae (Fig. 1B, “USA 2015, individual bees”) closely matched this single colony sample (Fig. 1B, “Virus-USA 2015”) with diversity profile Pearson’s correlation coefficients = 0.7453 - 0.9549 (S3 Table). Moreover, bees collected in 2010 in Texas and Maryland had DWV diversity profiles similar to the 2015 Maryland samples (Fig. 1B, “USA 2010, average”; 2010 and 2015 Pearson’s correlation coefficients = 0.5287 - 0.7432, S3 Table). Given the phylogenetic relatedness between the DWV consensus sequences from 2010 and 2015 (Fig. 3, S1 Fig., sequences labelled with suffix “-Cons”), a stable divergent DWV population has existed in *Varroa*-infested bees across the mainland US since at least 2010.

**Fig. 3.**
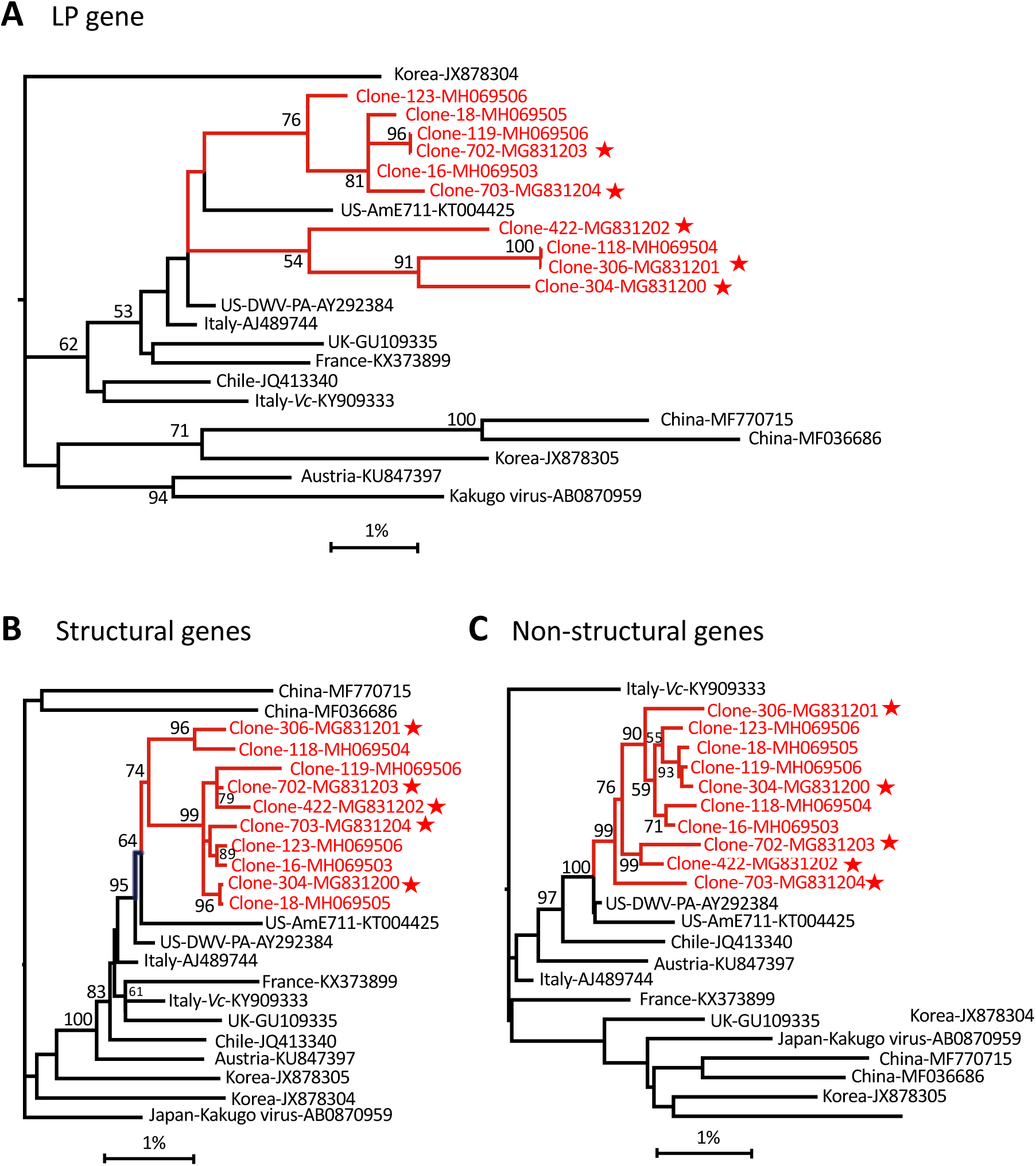
Phylogeny of genomic segments of DWV RNAs. Maximum likelihood phylogenetic trees were generated for the sequences coding for the (A) leader protein (LP; positions 1.1-1.7 kb), (B) structural proteins (positions 1.7-4.3 kb), and (C) major non-structural proteins (positions 4.3 - 10.1 kb). Nodes connecting the cloned sequences (“Clone-“), are shown in red, stars indicate DWV genotypes with tested infectivity. Bootstrap values for 1000 replicates are shown for groups with more than 50% bootstrap support. Scale shows genetic distance (%).

Phylogenetic analysis of the nearly full-length cloned DWV genomes (9.7 kb sections containing the full protein-coding sequences) produced for this study from a *Varroa*-infested Maryland colony revealed that they formed a cluster that also included the source colony’s NGS consensus (S1 Table, NGS library V99) and the consensus sequences from individual bees from the same apiary (S1 Fig., sequences labelled with prefix “Clone-”). This clade, which had 98% bootstrap support, was rooted in a branch with all complete DWV genomes from the US and the NGS-derived DWV sequences for 2010 and 2017 US bees from this study, which itself had 100% bootstrap support (S1 Fig). Specific genome sections revealed similar phylogenetic relationships (Fig. 3) further corroborating that the US DWV population was generated following a strong bottleneck event and subsequent diversification from a single, or closely related strain(s) from Europe.

### Design of the reverse genetic system for a virulent US DWV population

Stable co-existence of multiple variants within a virulent DWV population prompted questions about interactions between virus genotypes and driving forces maintaining high diversity. We used a molecular approach to investigate these interactions, which involved producing a series of full-length infectious cDNA clones of genomes of distinct isolates co-existing in a DWV population from a single colony (Fig. 3, S1 Fig., sequences labelled with a star). Together these clones captured a significant proportion of the genetic diversity present in a typical US DWV-A population (Fig. 1C, blue line). The colony used for virus preparation was *Varroa*-infested, contained honey bees showing wing deformities consistent with high DWV levels [2,5], and did not survive the following winter season, indicating that it harbored a virulent DWV strain typical of declining US colonies. The selected clones (n=5) and parental population (“USA-2015, 50 bees”) had highly similar distributions of divergent nucleotides (n=286) and amino acids (n=36) (Fig. 1C vs 1B). Shannon’s diversity profile for equal proportions of each clone (Fig. 1C, blue line) was positively correlated (Pearson R=0.804; S3 Table) with that of the parental DWV population (Fig. 1B, black line). When injected into honey bee pupae, full-length DWV *in vitro* RNA transcripts generated from the cDNA clones were infectious. The clone-derived DWV isolates replicated to similarly high levels, about 10^10^ to 10^11^ genome copies per bee, as wild-type RNA of the same DWV preparation used to design of the clones. Significantly lower (Wilcoxon rank sum test, P < 0.01) virus levels were observed in pupae injected with PBS, or with the mutant transcripts 304Δ and 306Δ, in which essential viral genes required for replication were deleted (Fig. 4A). Since DWV is widespread and present at low levels, about 10^5^-10^7^ copies per insect in virtually all honey bees including Varroa-free [12], this virus load was detected by qRT-PCR even in control samples. Therefore it was important to determine that the virus which replicated to high levels in the DWV transcript-injected honey bees was indeed clone-derived. The identity of clone-derived DWV progeny in the injected pupae was confirmed by both using the unique restriction site introduced into the clones (Fig, 4B) and by NGS analysis (S1 Table), which showed that the consensus nucleotides of the clone DWV sequences were identical to their respective parental cloned cDNA (Fig. 4C), while retaining low diversity (Fig. 1C, red graphs; Fig. 2, Group 5 – “USA 2015 cDNA clone-derived USA-2015”; S1 Table), thereby proving that the cloned DWV isolates were infectious.

**Fig. 4.**
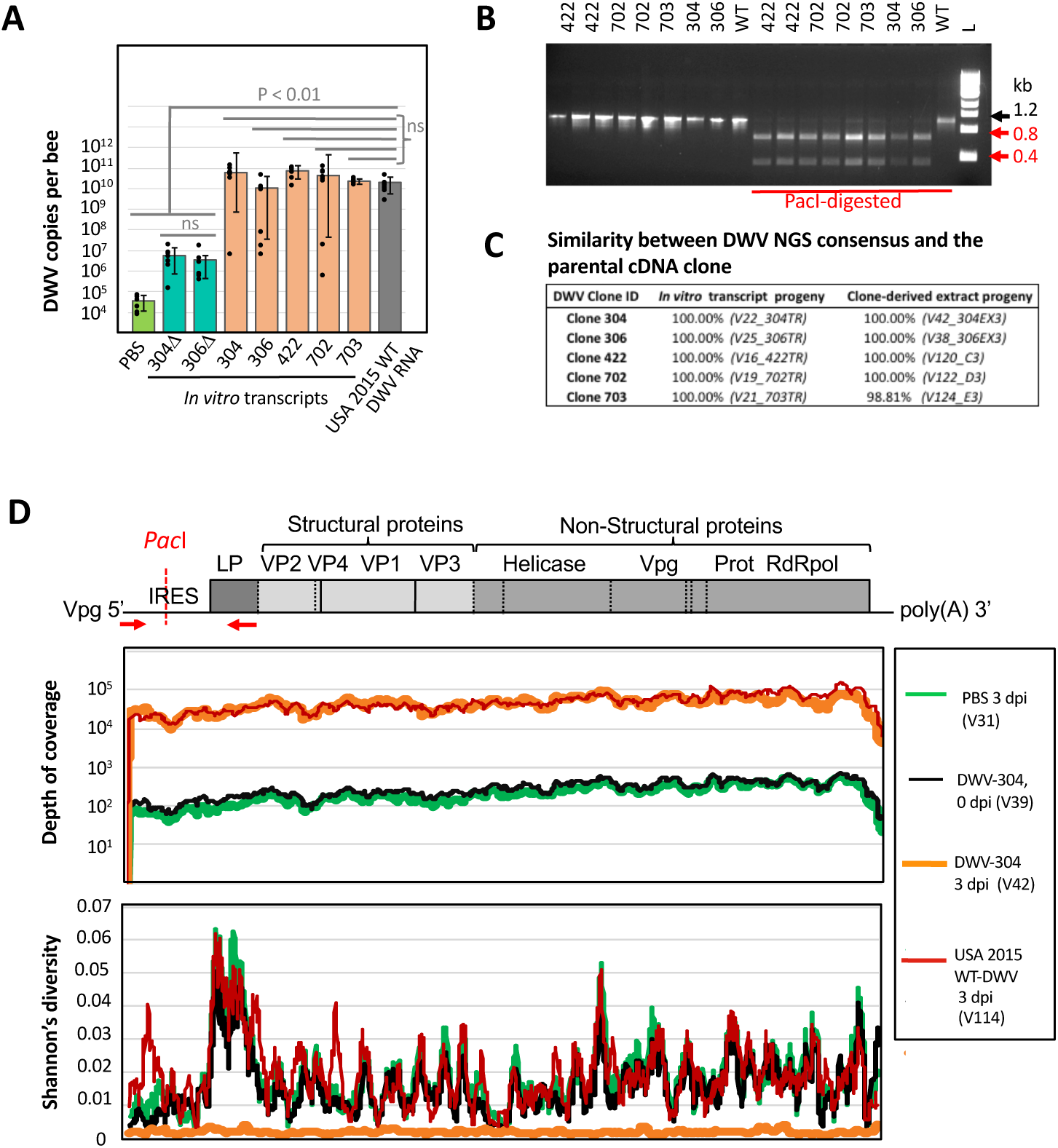
Infectivity of clone-derived DWV isolates. (A) Replication of DWV in honey bee pupae injected with 5 μg of *in vitro* RNA transcripts from full-length infectious DWV cDNA clones (304, 306, 422, 702 and 703), DWV cDNA constructs with a deletion of essential replication genes (Δ304, Δ306), 5 μg of wild-type DWV RNA (USA 2015 WT DWV RNA), or buffer control (PBS). The injected pupae, n=6 per each group, were sampled at 3 days post-injection (dpi), and the DWV loads per individual pupa were quantified by qRT-PCR. Black dots indicate DWV load in individual honey bees. The columns show the average DWV copy numbers for each treatment, ±SD. Statistically significant (P<0.01) and non-significant (ns) differences between treatments groups are indicated above the bars. (B) Presence of the introduced *Pac*I restriction site in the progeny of the clone-derived DWV genomes in the *in vitro* transcript-injected pupae at 3 dpi. (C) NGS analysis of the cDNA clone derived DWV progeny from the *in vitro* transcript-injected and clone-derived extract-injected honey bee pupae. Shown is the percentage of similarity between the DWV consensus sequences produced from NGS libraries (sample ID indicated in brackets) compared with their respective parental DWV cDNA sequences. (D) NGS analysis of the pupae infected with divergent wild-type DWV and clone-derived DWV (clone 304) immediately after injection (0 dpi) and at 3 dpi. NGS library identifier is contained within parentheses.

We then investigated replication dynamics of the recovered clone-derived virus isolates. After 48 and 72 hours, DWV levels were the same in pupae injected with 10^7^ genome copies of either clone-derived or wild-type DWV (Fig. 5A), further demonstrating that individual DWV genomes replicate to the same level as the parental divergent DWV population (Fig. 5A, Wilcoxon rank sum test, P > 0.01). Consensus DWV sequences at 3 days post-injection (dpi) were identical to their corresponding cDNA clones in all but one case, likely due to a background DWV infection in the injected pupae (Fig. 4C). DWV levels at 3 dpi were significantly higher (Fig. 5B, Wilcoxon rank sum test, P < 0.0001) than “0 dpi” pupae sampled immediately after injection with the virus, which had the same levels as PBS injected controls (Fig. 5B). The recipient honey bee pupae had low-titer background infection which caused DWV diversity to be high in “DWV-304 - 0 dpi” pupa injected with clone-derived DWV (Fig. 4D, black graphs - V39), but the clone-derived progeny had low, nearly clonal, genetic diversity at 3 dpi (Fig. 4D, orange graphs - V42), indicating that only the hemolymph-injected clone-derived virus replicated to high levels, rather than the highly divergent background, in stark contrast with pupae injected with wild-type DWV (Fig. 4D, red graphs - V114). Notably, in the honey bee pupa injected with buffer (PBS) control, divergent DWV remained at a low level (Fig. 4D, green graphs – V31). It could be argued that the mutation rate was too low to detect accumulation of mutant variants within 3 days post-infection, therefore future studies may wish to extend the post-injection analysis timeframe to allow for direct observation diversification of the clone-derived genotypes.

**Fig. 5.**
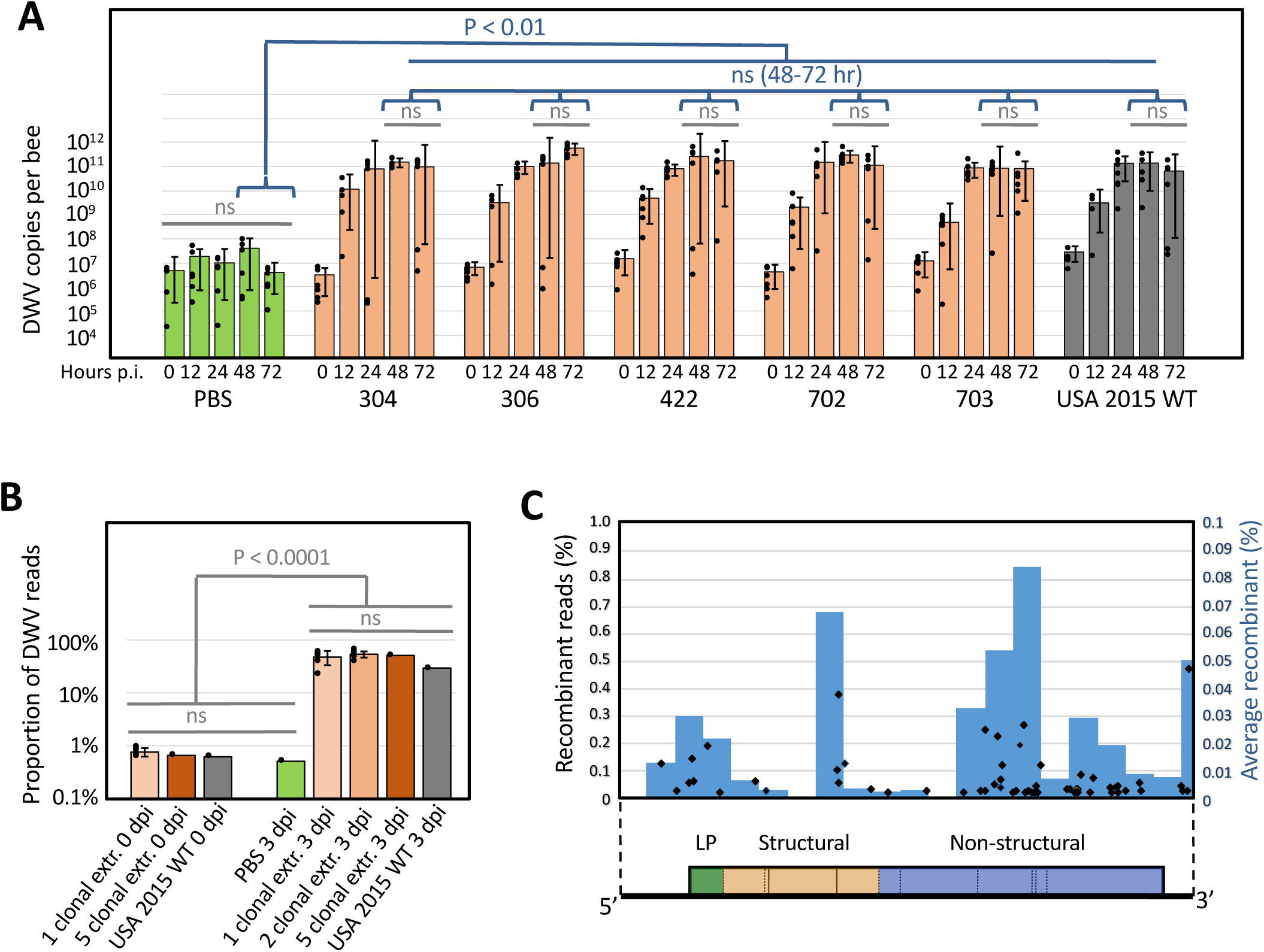
Replication dynamics and interaction between the clone-derived DWV isolates. (A) Average DWV RNA copies per honey bee pupa, ±SD. Individual pupae copy numbers indicated by black dots. (B) Proportion of DWV reads in NGS libraries for individual, mixed and control infections. Significant and non-significant (ns) differences are indicated. (C) Recombination breakpoints generated in mixed clone-derived infections. Black diamonds show proportion of recombinant reads in individual mixed infections (left Y axis). Blue bars show average proportion of recombinant reads in a genome section (right Y axis). DWV genetic map is shown below.

### Mixed infection of clone-derived DWV genotypes showed a lack of synergistic effects and competitive exclusion

Divergent virus populations might have increased fitness compared to individual genotypes due to complementation between genotypes that could reinforce each other to achieve a higher replication rate [18]. The replication dynamics of the individual clone-derived DWV isolates, their mixtures and divergent wild-type DWV were compared. When honey bees were infected by hemolymph injection with, in total, of 10^7^ particles of the clone-derived DWV isolates, there were no significant differences (Wilcoxon rank sum test) between DWV levels in pupae infected with either individual clone-derived isolates, all possible pair-wise combinations of these isolates, or a mixture composed of all five isolates (Fig. 5B). The DWV Shannon’s diversity profile in pupae injected with the mixture of all five clonal isolates was similar to that predicted assuming equal proportions of each component (Fig. 1B, black and blue lines, respectively; S3 Table, Pearson’s correlation coefficient P= 0.9825). Such a tight correlation between theoretical and observed profiles for mixed infections suggests each of five co-infected genotypes replicated to similar levels in the re-assembled population. NGS analysis of infections induced by pair-wise clone-derived isolate mixtures (S1 Table) allowed a genome-wide view of strain success *via* the ratios of alternate nucleotides at the expected divergent positions (S2 Fig.). Although some clones showed higher proportions than others, no complete competitive exclusion [19] was observed over 72 hours when resident genotypes of DWV where co-inoculated.

### Widespread recombination between the DWV isolates in mixed infections

NGS of the pair-wise clone-derived isolate mixtures also allowed us to investigate recombination within DWV populations. Analysis of the changes in the proportion of divergent nucleotides along the DWV genome (S2 Fig, lower panels, orange bars) revealed widespread generation of novel DWV variants as a result of recombination events between the clone-derived genotypes. This was confirmed by an analysis for structural variants that identified recombination breakpoints between clones (S2 Fig, upper panels). Recombination sites were clustered mostly in the regions preceding the leader protein (LP) region and in the LP region itself, the main 3’-proximal non-structural block in the helicase region, and the border of the regions coding for the structural VP1 and VP3 (Fig. 5C). Independent evidence for a high recombination frequency between DWV genotypes came from the phylogenetic analysis of the functional sections of the cloned full-length DWV genomes (Fig. 3). It showed different topologies in the phylogenetic trees, supported by high bootstrap values, for the genome sections coding for LP, the structural proteins and the non-structural proteins. For example, Clone-304 and Clone-18 have almost identical structural gene sections (Fig. 3B), but their LP gene sections are in different clades (Fig. 3A). Similarly the non-structural genes of Clone-119 and Clone-304 are very close (Fig. 3C), but their structural genes are in separate branches (Fig. 3B). Such differences in the tree topologies could be explained by reshuffling of genome sections between members of the DWV population.

### High genetic heterogenicity of DWV populations as a RNAi evasion mechanism

Why do wild-type DWV populations maintain such high genetic diversity in the absence of obvious synergetic effects, i.e. individual clones replicated to the same levels as mixtures of the clones and the parental divergent population? We suggest that diversification of DWV in honey bees might be driven by host RNA interference (RNAi) defenses and maintained as a way to evade sequence-specific RNAi similar to West Nile virus (WNV) in mosquitoes and Powassan virus in ticks [20–22]. These studies showed that rare genetic variants of these viruses avoid control by RNAi targeting major variants due to the lack of complementary between the minor strains and the guiding small interfering RNAs (siRNAs) derived from the major viral strains [20–22]. Analysis of the distribution of polymorphic sites in DWV populations provided several findings in support of this suggestion.

We found that most of the nucleotide changes in sampled DWV populations were silent, with a mean ratio of synonymous to nonsynonymous substitutions (dS/dN) in the tested infectious clones reaching 24.7, indicating strong purifying selection (S4 Table). Accumulation of a high number of nucleotide changes throughout DWV genomes, constrained by the need to maintain the same coding capacity, suggests that the virus is forced to explore sequence space to maintain its high diversity. This is consistent with the hypothesis that diversification of DWV RNA is driven mostly by selection of novel sequence variants of DWV RNA capable of evading specific RNAi targeting due to mismatches with siRNAs. Analysis of the NGS libraries confirmed that polymorphic sites are present throughout DWV genomes (Fig. 1B), in agreement with a previous report that the entire DWV genome could be targeted by RNAi [12] and therefore subjected to diversification. Still, a significantly higher density of polymorphic positions was consistently observed in the LP region and in the 3’ proximal part of DWV coding for the non-structural block (S3 Fig.; S4 Table). Additionally, the cloned infectious DWV isolates showed a subset of codons (n=31) with dS/dN<1, which are potentially subjected to diversifying positive and/or relaxed selection. Interestingly, 12 of these putative positively selected codons were located in the LP region, a ten-fold higher density than in the rest of the genome (S3 Fig., S4 Table).

To estimate the potential of US DWV populations to evade specific RNAi established to a single given DWV strain, we analyzed DWV NGS libraries from a typical colony, a single honey bee sampled in 2015, and a pool of 8 honey bees sampled in 2010 (S1 Table; Libraries B2_PBS, V57, and V99) for sequence variants capable of avoiding specific RNAi targeting, by recording the number of alternate nucleotides (occurring above 1% and 10% levels of the read coverage) in a sliding 22-nt window (Fig. 6), i.e. the size of a potential siRNA target of in the honey bee [12]. Within the colony-level DWV population, most of the genome, 81.80%, showed at least one sequence variant present at a frequency of 1% or higher, and 38.81% of the genome showed variants present at 10% or higher. Notably, the polymorphic 22 nt windows covered a significant proportion of the viral genome in the DWV populations from the individual bee sampled in 2015, with 35.53% and 41.31% of the genome with sequence variants present at 1% or higher and 10% or higher, respectively. Similarly, the DWV population from a pool of 2010 US honey bees showed 57.42% and 33.01% divergent coverage. Clearly, divergent DWV populations harbors a pool of readily available genetic variants some of which may possess reduced vulnerability to specific RNAi. Therefore, we propose that the widespread recombination events in DWV populations between divergent co-existing DWV genotypes contributes to the generation of variants less effectively targeted by established RNAi and allow DWV to evade this sequence-specific antivirus response.

**Fig. 6.**
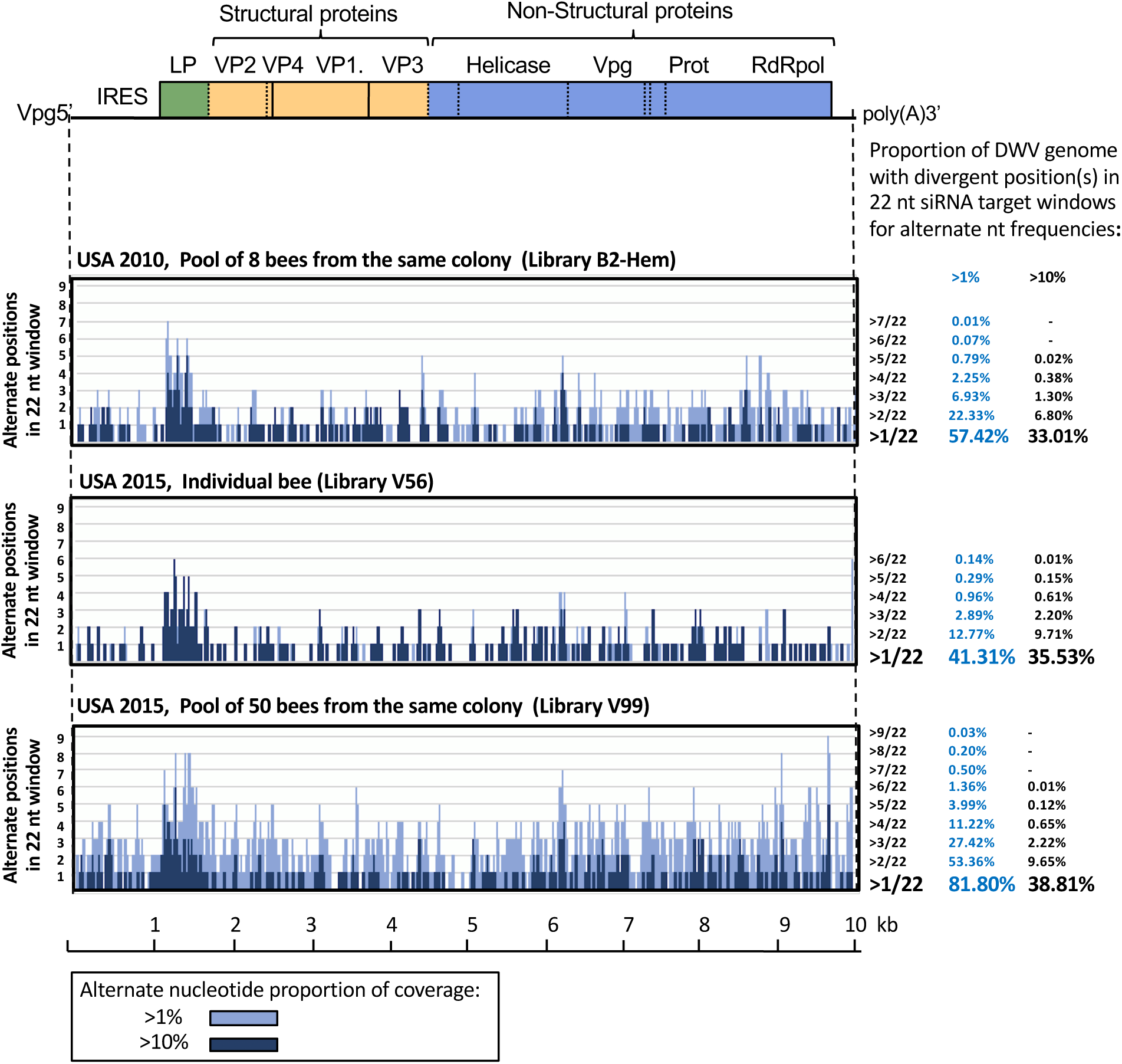
Potential contribution of DWV population diversity to evasion of sequence-specific RNAi targeting. Distribution of the number of alternate positions in 22 nt -long potential siRNA target windows for the polymorphic sites with more than 1% (light blue graphs) and 10% (dark blue graphs) of NGS coverage. DWV genetic map is shown above the graphs. Summaries of the DWV genome coverage (%) with different number of mismatches in 22 nt sliding window for each of the NGS libraries is shown at the right of the graph.

### Natural differences between DWV genotypes might significantly affect efficiency of antiviral RNAi targeting

We sought to experimentally test if natural variation between DWV genotypes co-existing in a single population was sufficient to significantly impact the effectiveness of antiviral RNAi and indeed could act as a RNAi evasion mechanism. It was shown previously that antiviral RNAi in honey bees could be induced by introduction of dsRNA molecules orally [23]. To do this, we produced double-stranded (ds) RNA specific to two co-existing isolates US DWV-304 and DWV-422. The targeted 283 nt region of DWV genomic RNA (positions 1242 to 1524 of the GenBank accession MG831200) had 5.3% divergent nucleotides between isolates DWV-304 and DWV-422, which mostly resulted in one mismatch in any given 22 nt sliding window (Fig. 7A). In two independent experiments (design summarized in Fig. 7D) newly emerged adult honey bees from colonies with low *Varroa*-infestation levels were fed with 1 ⍰g of the dsRNA (ds304 or ds422) in 5 ⍰L of 50% sucrose either with or without the virus extract containing 10^8^ particles of the clone-derived DWV-304 or DWV-422. The bees were maintained for 7 days prior to sampling and quantification of DWV loads in individual insects. The identity of clone-derived DWV variants was confirmed by sequencing the RT-PCR fragments corresponding to the LP region.

**Fig. 7.**
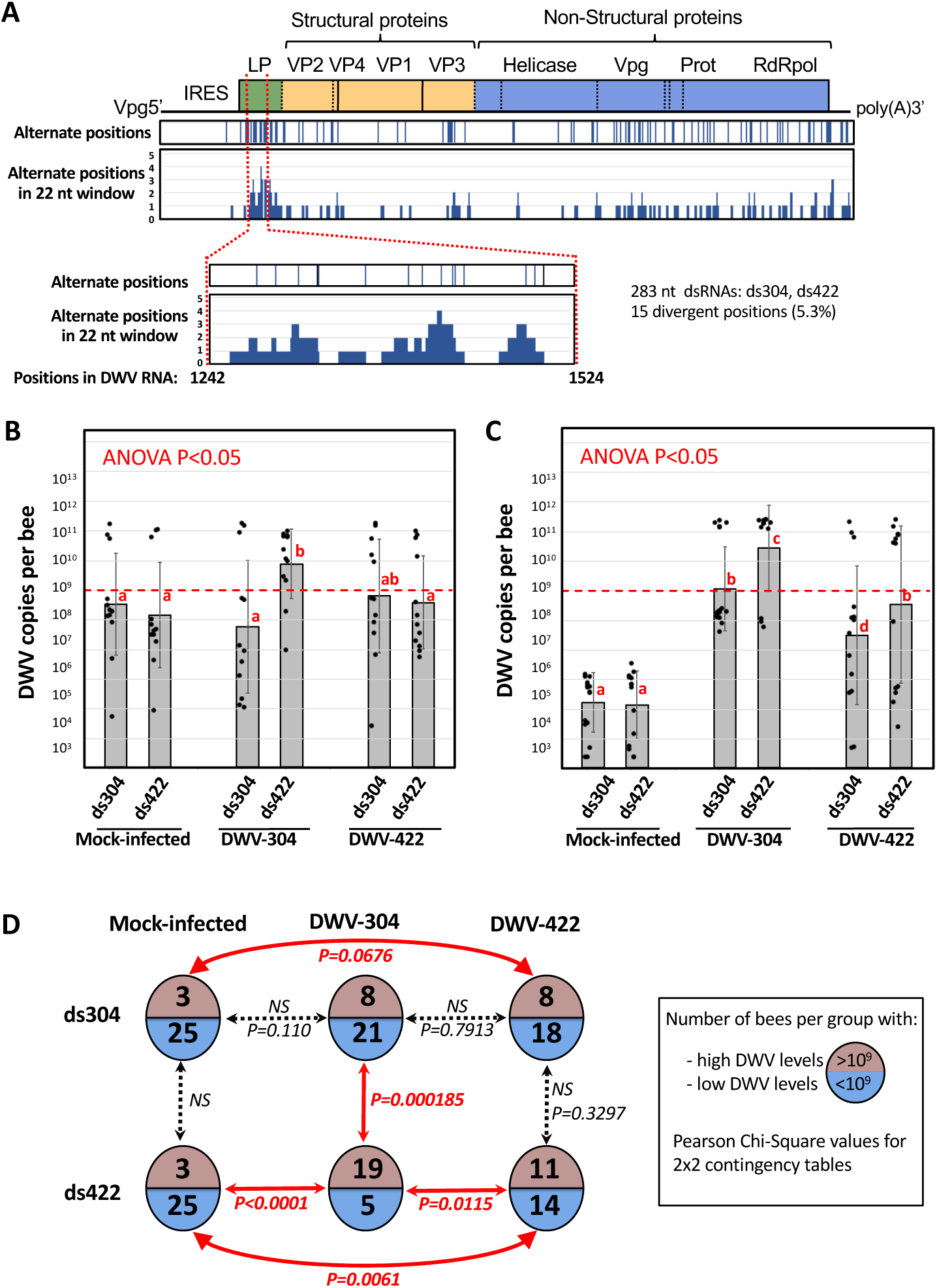
Impact of specificity of RNAi targeting on outcome of DWV infections. (A) Design of the 283 nt dsRNA specific to the DWV isolates 304 and 422(ds304 and ds422) corresponding to the regions 1242-1524 of DWV genome (GenBank Accession MG831200). (B, C) DWV loads in the individual adult honey bees 7 days post feeding with 10^8^ clone-derived virus particles and 1 ⍰g of the DWV-304- or DWV-422-specific dsRNA. Columns show average copy number per treatment groups, ±SD; black dots indicate DWV copy numbers in individual honey bees. The groups with significantly different (P<0.01, ANOVA) DWV copy numbers are indicated above the bars with different letters. The red line indicate threshold level of 10^9^ DWV genome copies per honey bee which was used to categorise DWV levels as low or high. (C) Analysis of the DWV infection outcomes in two oral inoculation experiments with the strain-specific dsRNAs. Circles indicate experimental groups, numbers of the bees in each groups with high (> 10^9^ copies) or low DWV (below 10^9^ DWV copies) levels of DWV are shown on the top or bottom of the circles. A 2×2 contingency table analysis was used to test significance of the differences in proportions of the bees with low or high DWV levels.

The outcome of DWV-304 infection was significantly influenced by the specificity of the dsRNA. There were significantly higher levels of DWV in the DWV-304-infected bees which received ds422 (Fig. 7B,C; Group DWV304/ds422) compared to the group where this virus strain received ds304 (Fig. 7B,C; Group DWV-304/ds304), p<0.05, ANOVA. The virus levels in individual bees showed bimodal distribution, which allowed us to categorize DWV loads as low or high, with a 10^9^ copies threshold, and carry out a 2×2 contingency table analysis. We found, in the case of the bees infected with DWV-304, a significantly reduced number of bees developed a high-level DWV infection in the group “DWV-304/ds304” (which received ds 304 perfectly matching virus isolate) compared to the group “DWV-304/ds422” (which received ds422 mismatching the inoculated virus), P=0.000185 (Fig. 7D). Notably, the proportion of bees with high DWV levels in the group “DWV-304/ds304” were not significantly different from that in control group “Mock-infected/ds304” (Fig. 7D). Similarly, ds422 suppressed development of high DWV levels more efficiently in the case of inoculation with DWV-422 compared to DWV-304, P=0.0115 (Fig. 7D, Groups “DWV-304/ds422” and “DWV-422/ds422”). These results suggest that natural variation between DWV genotypes within a single population (e.g. DWV-304 and DWV-422) might significantly reduce the efficiency of RNAi targeting.

## Discussion

Analysis of DWV population genetics may provide novel insights into the factors that contribute to sharp increases in viral pathogenicity. In the case of DWV viral populations with different degrees of virulence are available and the emergence of highly virulent DWV has been linked to the spread of the mite *Varroa destructor* [5]. *Varroa* which appeared in the US in 1987 [24] and is currently detected across the US in 84.9% of migratory beekeeping operators and 97.0% of stationary apiaries [25]. While DWV is transmitted through vertical and oral routes [2,9], the mite provides an efficient route for horizontal transmission of DWV *via* direct injection into the honey bee hemolymph. Through *Varroa*-mediated transmission, survival and propagation of DWV became less dependent on the survival of infected bees thereby selecting viral variants with increased replication levels. It is unclear which genetic changes are associated with *Varroa*-mediated transition, but selection of certain strains of the DWV may be involved, such as recombinants between DWV-A and VDV1 [11–13]. Additionally, reduction of genetic diversity in *Varroa*-transmitted DWV to nearly clonal virus populations was reported in two independent studies of the honey bees from the UK in 2013 [12] and the Hawaiian islands in 2009 [9] in stark contrast to the diversity of DWV present in the covertly infected, honey bees with low DWV levels [9, 12]. Surprisingly, in this study we found that low genetic diversity of DWV is not an universal feature of *Varroa*-associated DWV. By using NGS we demonstrate that the virulent DWV currently circulating in the US *Varroa*-infested honey bee colonies has significantly higher diversity than the DWV in the UK 2013 study [12], which used the same high-throughput sequencing approach (Fig. 2A). Diversity of DWV was equally high in individual US bees regardless of virus load (Fig. 2, Groups 3 and 4), while bees sampled in the UK in 2013 had a significantly lower virus diversity in the bees with high DWV loads (Fig. 2 A, Groups 1 and 2).

Differences between UK and US DWV genetic diversity levels are dramatic when colony-level virus populations from pools of 50 bees are compared (Fig. 2A,B). Genetic diversity of the UK and the US colony level-samples (50 bees) were not singificantly different from individual bees with high DWV levels in their respective locations (Fig. 2A, UK 2013: P=0.7408, US 2015: P=0.326, ANOVA). This suggests that while the individual bees sampled in the UK were infected with the same or very similar DWV variants, while individuals in the US colony were infected with a many divergent genetic variants of the virus. Indieed, simiular distribution profiles of polymorphic postions were observed in the NGS libraries of individual bees and a colony level population in sampled in the US (Fig. 1B, S3 Table) suggesting co-existence of the same multiple DWV genotypes even in individual US bees. Phylogenetic analysis of the full-length coding sequences of DWV genotypes from a single colony-level population sampled in 2015 showed monophyletic origin of co-existing DWV isolates (S1 Fig., bootstrap value of 98%) and the genomic sections (Fig. 3), indicating that this diversity was generated as a result of diversification of a single parental DWV genotype. Therefore, although samples from a period in time when low DWV diversity occurred in the mainland US have not been analyzed, there exists supporting evidence that a bottleneck did occur sometime prior to 2010 when high diversity was measured (Fig 1B, grey graph).

The degree of divergence between some genotypes, within a given virulent US DWV population, infecting individual insects exceeds that reported for DWV isolates sampled in different locations several years apart. For example, co-existing US DWV isolates 703 and 306 differ from each other as much as the North American DWV-PA reported in 2006 [1] and DWV-*Vc*-Italy discovered in the *Vespa crabro* wasp in 2017 [26] (S1 Fig.). Such high heterogeneity in DWV populations highlights the importance of comprehensive characterization of the RNA virus populations, and therefore high heterogeneity should be considered when evolutionary histories of this virus are modelled based on phylogeography especially as sampling is limited in many locations [10]. US DWV has significantly higher diversity when compared to other RNA viruses, including those infecting invertebrates. For example, an analysis of the DWV cDNA clones from a typical individual colony, with a degree of diversity similar to that in individual honey bees (Fig. 1B) had a 1.26% mean nucleotide diversity between the five infectious cloned isolates, whereas an assessment of the cloned West Nile virus fragments in the wild-collected mosquitoes and birds showed a mean interhost diversity of only 0.237%. [27].

Further investigation of interactions between DWV genotypes within the virulent DWV-A population in a typical declining *Varroa*-infested US colony required isolation of individual components of the population. Due to the absence of DWV-free honey bee tissue culture for DWV [16], essential for isolation of the individual components of virus populations by “classical” plaque assay [28], we used a molecular approach to obtain DWV isolates. This involved the design of a series of infectious cDNA clones corresponding to individual RNA genomes forming a DWV population. Notably, the entire protein coding sections of the DWV RNA in the cDNA clones derived from the existing individual RNA molecules by using nearly full-genome amplification rather than being assembled, thereby constructing a “snapshot” of the quasispecies. In doing so, we effectively designed a reverse genetic system for an invertebrate RNA virus “mutant cloud” [29], this being the first of its kind.

One of the key drivers for diversification of the DWV population could be complementation between genotypes in mixed infections as reported for other RNA viruses [18]. To test this hypothesis we compared replication dynamics of individual isolates and the parental DWV population, combinations of the clones, and parental wild-type DWV by in pupal injection test. Surprisingly, we established that 3 days after injection the levels of virus were not significantly different in all cases, except the buffer (PBS) control (Fig. 5B). In addition, the individual clone-derived isolates and wild type DWV showed the same replication dynamics, and were replicating to similarly high levels (Fig, 4A, Fig. 5A). Taking together this suggested that no complementation between genotypes of DWV population took place. At the same time co-infection experiments with cloned isolates demonstrate their mutual compatibility and the lack of any strong competitive exclusion phenomena (S2 Fig.). It should be noted, that such co-existence of the divergent DWV isolates is not a rule, for example in the mixed experiment when DWV populations from different UK locations were co-injected, DWV isolate derived from the *Varroa*-infested bees over-competed that sourced from *Varroa*-free region [12]. It is therefore very likely that US DWV isolates existing in the same population were adapted to each other in order to maintain high population diversity. We also established that recombination events between the DWV isolates in a population were widespread, as evidenced by different topologies of the phylogenetic trees for genome sections of the full-length clones (Fig. 3) and directly detected recombination events as observed in the virus progeny in the case of mixed infections of cloned isolates (Fig. 5C; S2 Fig.).

These results prompted questions on what drives diversification of DWV and the maintenance of high diversity in the absence of apparent complementation between isolates, and, in addition, how the contradiction between high DWV diversity in the Varroa-infested US DWV and the previously reported nearly clonal nature of the *Varroa*-associated DWV in the UK (12] and Hawaii [9] could be explained. We therefore proposed the following model of DWV diversity dynamics. Firstly, the introduction of *Varroa* reduced DWV diversity through a selective sweep as a result of selection of the virus variants capable of being transmitted by *Varroa* mites from diverse pre-*Varroa* population, as seen in bees from the Hawaiian island Oahu in 2009 [9]. Although it cannot be completely ruled out that *Varroa*-associated DWV variants may be introduced with the mite itself, this scenario is unlikely because partial viral RNA sequencing demonstrated that the *Varroa-*selected variant was present in pre-*Varroa* divergent DWV [9]. Notably, this model does not indicate that DWV variants selected as a result of DWV transmission have to replicate in *Varroa* mites, but might instead may have better stability in *Varroa* mites and/or better replication when injected directly into the honey bee hemolymph. Following this selective sweep, a period of *Varroa*-adapted DWV strain diversification *via* point mutations and recombination was undertaken. We suggest that the main driver for such diversification is the generation of DWV population, one capable to circumvent the genotype-specific antiviral defenses, and in particular RNAi responses. This model could explain the co-existence and maintenance of multiple genetic variants. Indeed, distribution of the divergent positions in the DWV genome suggests that the virus explores sequence space to maximize the proportion of genome containing mismatches in potential 22 nt siRNA targets, enabling it to evade specific RNAi (Fig. 6). We further demonstrated that the natural degree of variation observed between clones co-existing is a single population (e.g. clones 304 and 422) could result in such evasion of the strain-specific RNAi (Fig. 7).

The presence of alternate DWV variants potentially capable of evading specific RNAi, which may cover around 80% of the genome in a colony-level population for >1 % alternate level (Fig. 6), potentially enabling the virus population to quickly respond to specific RNAi by increasing the proportions of less-efficiently targeted genotypes with mismatches in the targeted region (Fig. 6). In addition, a high recombination rate observed in the DWV populations (Fig. 5C, S2 Fig.) could result in the assembly of novel genotypes composed of genome sections less efficiently targeted by a given set of siRNAs, thereby enabling RNAi evasion. Most notably, this scenario requires the ability of individual genotypes to be independent from each other, this is consistent with the equal replication rates of isolates (Figs. 4A and 5A) and the lack of complementation as observed in the mixed infections (Fig. 5B). In other words, maintaining a high genetic diversity is advantageous for the virus as the because recombining and changing proportions of the existing variants occur much faster than the accumulation of mutations, thereby enabling an instantly changing composition of predominant genotypes in response to the different pressures, including RNAi.

This model of DWV population dynamics, along with both previous and current findings, fully supports the view that mutability of RNA viruses is essential in maintaining virulent phenotypes [30]. A selective sweep of DWV following the introduction of the novel mite vector favours the subsequent generation of novel *Varroa*-adapted variants with superior virulence and capacity to escape antiviral defenses (e.g. being targeting by specific RNAi established against predominant genotypes as shown for other invertebrate RNA viruses [20–22]) in a process similar to “punctuated immune escape” [31], which was previously proposed for both vesicular stomatitis virus [32] and Influenza virus A [33,34].

A rapid expansion of one of many low frequency genetic variants of a virus, which occurs due to increased fitness, results in the temporary reduction of overall virus population diversity [31]. Indeed, very low diversity of DWV in the UK colony level sample (Fig. 2A, B, “Colony 50 bees, UK-2013”) suggests that this population was sampled at the point of sharp reduction. Subsequently, DWV diversity reverts back to high levels through accumulation of mutations, ultimately establishing virus populations with multiple isolates that co-exist even in individual insects (Fig. 2A, Group 4), with a high colony level diversity (Fig. 2A, B, “Colony 50 bees, USA-2015”, pointed with green arrow). Indeed, our initial analysis of DWV in the 2015 samples from Oahu Island showed a high variability that matched that currently observed in the mainland US (S4 Fig., S5 Fig.). Thus, such a long-term, semi-stable coexistence of virus isolates may arise spontaneously in virus populations as observed in bacteria [35]. It is notable that our phylogenetic trees along with our NGS evidence from 2010 suggest that the US mainland bottleneck in DWV diversity occurred before 2010 and that the introduction of *Varroa* to the US mainland occurred in 1980s [10,24]. Further analysis of additional samples collected shortly before and after the US mainland *Varroa* introduction, and/or samples from Hawaii prior to the *Varroa* introduction, would serve to confirm the role of *Varroa* in the selective sweep and significantly reinforce our model.

Our study of DWV has provided new insights into the divergent DWV population structure, including interactions between the viral genotypes within invertebrate RNA virus quasispecies and virus diversity dynamics by using a reverse genetic system and high throughput sequencing approaches. The high genetic diversity of DWV and its potential to instantly respond to the pressures of sequence-specific RNA-mediated antiviral defense, should be taken into account for the further development of anti-viral interventions, including those based on RNAi [23] or superinfection exclusion [36] mechanism, all of which may be used to improve honey bee health and ensure sustainable food security [8]. DWV is a member of the family *Iflaviridae*, [37], which infects species of different insect taxa [38], therefore further analysis of the DWV gene functions and its interaction with the host, using our reverse genetic system as developed in this study, could be applied to other viruses and therefore contribute to the further understanding of fundamental aspects of RNA virus infections and evolution in invertebrates and their wider ecological impact.

## Materials and Methods

### Honey bees

*Varroa*-infested honey bee colonies maintained by the USDA-ARS Bee Research Laboratory (Beltsville, Maryland, USA), which had high *Varroa* infestation (15-20% of pupae infested with mites) and bees showing wing deformities consistent with high DWV levels, were used as the source of mite-exposed honey bee pupae for NGS analysis and isolation of DWV particles in October, 2015. Unparasitized honey bee pupae from colonies with low *Varroa* infestation (with below 1 % of mite-parasitized pupae) were used in the injection experiments. Newly emerged pupae from the Maryland colonies with low Varroa infestation were used in the oral infection experiments. A colony from Texas was used in the *in vitro* RNA transcript and wild-type RNA injection experiment. A colony from Florida was used in the experiment involving injection of virus extracts, both clone-derived and wild-type.

### Sequence analysis

The NGS libraries for this study were produced by Illumina HiSeq2500, with each containing 12 to 14 million paired-end reads (S1 Table; NCBI BioProject PRJNA431793, Sequence Read Archive study SRP135682). These libraries, along with the previously published UK NGS libraries [12] (S2 Table 2), were analysed as described previously [17]. Briefly, Illumina reads were trimmed to remove adapters, contaminant sequences and low quality bases. Cleaned data from each sample library were individually aligned to reference sequences of three distinct strains of DWV-like viruses, DWV-A (GenBank Accession GU109335), VDV1 (GenBank accession AY251269) and DWV-C [4] using Bowtie2. Nucleotide counts, coverage and Shannon’s diversity index estimates for each nucleotide position were calculated from SAMtools mpileup output. Although Illumina HiSeq technology has a very low per-nucleotide substitution error rate (0.08% and 0.12% for the first and second reads, correspondingly [39]), to reduce sequencing chemistry errors, the Shannon’s diversity index estimate was further corrected for potential errors using approach described previously [40]. In brief, the error model was calibrated using data from an Illumina library from a bee injected with a cloned virus of known sequence. Data from positions 5230 to 6600 of the DWV genomic RNA, representing a low diversity region of the virus, were used to calculate an alphahat of 9.88*10^-5^. To quantitatively assess the degree of similarity between different DWV genetic diversity profiles, we calculated the Pearson correlation between the Shannon’s diversity index profiles averaged for 100 nt windows. For detection of recombination breakpoints in the honey bee pupae injected with the mixtures of the cloned-derived DWV genotypes, cleaned data from each sample NGS library were individually aligned to the VDV1 and DWV references using SpeedSeq (version 0.1.2). Structural variants were detected using the lumpy smooth script from the LUMPY package (version 0.2.13) and additional genotype and coverage metrics for each variant were calculated using SVTyper, a script within the SpeedSeq package. Breakpoint positions representing a recombination between two different viral isolate sequences were considered for further analysis if they had evidence from more than 10 supporting events (split or discordant reads) and exceeded 0.02% of NGS coverage (the level was chosen to ensure that the reads corresponded to viable, i.e. replicating, recombinant genomes, rather than aberrant variants). Phylogenetic analysis was carried out by aligning the viral nucleotide sequences using CLUSTAL and producing maximum likelihood and neighbour-joining trees, which were bootstrapped using 1000 replicates with the RAcML and PHYLIP packages, respectively. Synonymous (dS) and non-synonymous (dN) variations in viral genomes were identified using all known full-length North and South American DWV-A sequences (S1 Fig.) using tools provided by the Los Alamos National Laboratory’s HIV Databases (https://www.hiv.lanl.gov [41]); putative positive diversifying selection was assumed when dS/dN < 1 (S4 Table).

Analysis of DWV diversity in Hawaiian (Oahu island) apiary-level samples collected in 2015 as a part of the USDA-APHIS honey bee survey [24] included RT-qPCR amplification of the DWV cDNA fragments corresponding to the LP and the RdRp regions, using the oligonucleotide primers “DWV-LP-For” and “DWV-LP-Rev”, “DWV-POL-For” and “DWV-POL-Rev” (S5 Table), respectively, and direct sequencing of the RT-PCR products to identify polymorphic sites in the electropherograms. The RT-PCR fragments corresponding to the LP region were also cloned into a pGem-T easy vector (Promega) and Sanger sequenced.

### DWV cDNA clones and clone-derived viruses

Viral particles were purified by caesium chloride gradient centrifugation from a pool of 50 pupae out of a single *Varroa*-infested colony at the Bee Research Lab in Beltsville, Maryland, USA in October 2015 as described previously [11]. Viral RNA was extracted from the virus preparation using the RNeasy kit (Qiagen). The first cDNA strand corresponding to the entire 10 kb-long genomic RNA was produced using Superscript III Reverse transcriptase (Invitrogen) and the oligonucleotide primer “DWV-PmeI-A27Rev” complementary to the 3’ terminus of the DWV genomic RNA, which was preceded by the 27 nt oligo dT and the *Pme*I restriction site (S5 Table). The cDNA was used as a template to amplify a nearly full-length, 9.7 kb DWV RT-PCR fragment containing the full protein-coding sequence using the oligonucleotide primers “DWV-PmeI-A27Rev” and “DWV-USA-PacI462F”, identical to positions 472-502, preceded by the *Pac*I restriction site, (S5 Table) and the proof-reading thermostable DNA polymerase Phusion (New England Biolabs). The library was cloned into the pCR4Blunt-TOPO vector (Invitrogen) according to the manufacturer’s instructions. Ten clones containing 9.7 kb viral cDNA inserts were Sanger sequenced (GenBank Accession numbers MG831200 - MG831204, and MH069503 - MH069507). A set of five divergent DWV cDNA constructs (GenBank Accession numbers MG831200 - MG831204) was selected to construct a set of full-length infectious DWV cDNA clones capturing significant proportion of colony-level diversity. This was accomplished by inserting at the position corresponding to the 5’ part of the DWV cDNA the synthetic fragment “T7-Ribo-5’-DWV” (S5 Table) containing the T7 RNA polymerase, the 5’ ribozyme sequence [42] and the 462 nt-long 5’ terminal part of the DWV genome, using the *Not*I and *Pac*I restriction sites to create a series of 5 plasmids with different full-length cDNA inserts. The 462 nt-long 5’ terminal sequence was designed according to the DWV USA NGS data obtained in this study and included an extended 5’ terminal sequence (as previously [43] and in the GenBank Accession number KT215904). To produce control constructs p304Δ and p306Δ containing non-infectious DWV cDNA, the regions 7128-7998 and 7128-8347 coding for the viral protease region were deleted from clones p304 and p306 (GenBank Accession numbers MG831200 and MG831201), respectively, creating a frame-shift in the viral ORF that abolished translation of the viral RNA-dependent RNA polymerase. Plasmid constructs were linearized by using the *Pme*I restriction site located at the 3’ end of DWV cDNA, downstream of the A27 sequence, to produce the templates for a full-length kb transcript. The *in vitro* transcripts were produced using HiScribe T7 High Yield RNA Synthesis Kit (New England Biolabs) according to the manufacturer’s instructions. After 3 hr incubation at +37°C, the DNA templates were digested using TURBO DNase (Life Technologies). The RNA transcripts were purified using the RNeasy mini kit (QIAGEN), eluted with RNAse-free sterile water and stored at −80°C prior to use.

RNA transcripts were injected into honey bee pupae without *Varroa* mite feeding exposure. These pupae were extracted from brood frames with low *Varroa* infestation (approximately 1 mite-infested cell per 500) from a Texas apiary in December 2016. The pupae at pink eye stage were injected intra-abdominally into the hemolymph with 5 micrograms (9.3*10^11^ copies) of the *in vitro* transcripts or wild type DWV RNA in 10 ⍰L of phosphate-buffered saline (PBS), or 10 ⍰L of PBS control using syringes with a 0.3 mm needle G31 (BD Micro-Fine), essentially as described previously [12], and were incubated at +33°C, relative humidity 80% for 3 days to allow replication of the virus prior to RNA extraction and sequencing. It was necessary to inject high copy numbers of naked viral RNA to reliably intitiate clone-derived infection because, as expected, infectivity of naked RNA was very low compared to an encapsidated RNA. Notably, the injected RNA, either *in vitro* transcript or extracted from virus particles, were almost entirely degraded in the hemolymph after 3 days, qRT-PCR quantification of DWV in the pupae injected with 9.3*10^12^ copies of the control non-replicating DWV *in vitro* transcripts p304Δ and p306Δ had between 1.3*10^5^ to 1.7*10^7^ genome copies detected (Fig. 3A). For comprehensive characterization of the transcript-derived progeny, NGS libraries were generated from individual pupae each containing 12 to 17 million paired-end 150 nt reads with the share of DWV reads ranging between 43% to 63% (S1 Table 1).

For preparation of the clonal DWV extracts containing infectious DWV virus particles, individual transcript-injected pupae were homogenized with 2 mL of PBS at 3 days post injection. For each individual pupal extract, 1 mL was used for preparation of the DWV inocula, which included clarification by centrifugation at 3000g for 5 min and filtration through 0.22 ⍰m nylon filter (Millipore). The DWV concentration in the extracts was quantified by RT-qPCR, and the extracts were stored at −80°C prior to use. To confirm the identity of the clonal DWV in the extracts with their respective parental DWV cDNA clones, the DWV RNA region from positions 100 to 1300 nt was amplified by RT-PCR, sequenced and also digested with *Pac*I restriction enzyme.

Pink eye stage honey bee pupae, with no exposure to *Varroa* mite feeding, were extracted from a Florida brood frame with low *Varroa* infestation (1 infested cell per 500) in January 2017. Pupae were intra-abdominally injected into the hemolymph to introduce either the filtered DWV extracts (10^7^ genome copies in 10 ⍰L of PBS), which included the clone-derived recovered DWV variants that originated from transcript-injected pupae, the wild-type DWV, or buffer control (PBS). We sampled time series from “Time 0” (immediately after injection) through to 3 days post injection, and the DWV RNA copy numbers were quantified in individual bees by RT-qPCR as described previously [17].

### Assessment of the effect of strain-specific dsRNA on DWV replication

Double stranded (ds) RNA specific to the DWV clones were produced *in vitro* with T7 RNA polymerase (HiScribe T7 RNA polymerase, New England Biolabs) using as the templates PCR fragments corresponding to the positions 1242-1524 of the clones DWV-304 and DWV-422 (GenBank accession numbers MG831200 and MG831202, respectively) with T7 promoter sequences at the both 5’ and 3’ termini. These PCR fragments were amplified using the US DWV cDNA clones p304 and p422 as the templates and the primers shown in S5 Table. Oral infection included feeding individual newly emerged bees with 5 ⍰L of 50% sucrose containing 1 ⍰g of purified dsRNA (ds304 or ds422) and 10^8^ clone-derived virus particles, DWV-304 or DWV-422, or no virus in the case of control groups (Fig, 7D). The honey bees sourced from Maryland colonies with low Varroa infestation levels (less than 1% of pupae being infested with the Varroa mites) and were starved for additional 3 hours before feeding. The bees were kept for 7 days essentially as described in [44] in cages at +33°, 75% relative humidity, with *ad libitum* sources of water and 1:1 sucrose/water syrup w/v, prior to RNA extraction,

### Data deposition

Raw sequence data and consensus DWV sequences have been deposited at DDBJ/EMBL/GenBank within BioProject PRJNA431793 under the Sequence Read Archive (SRA) study SRP135682 (accessions SRR6833910 - SRR6833954) and Transcriptome Shotgun Assembly (TSA) study GGSE00000000 (accessions GGSE01000001-GGSE01000045), respectively. Cloned viral cDNA sequences are available in GenBank with accession numbers MG831200 - MG831204, MH069503 - MH069507, and MH594118 - MH594121. The DWV cDNA plasmids are available for research purposes from USDA Bee Research Laboratory upon request.

### Note on not endorsing products mentioned in the methods

Mention of trade names or commercial products in this publication is solely for the purpose of providing specific information and does not imply recommendation or endorsement by the U.S. Department of Agriculture.

## Acknowledgements

This research was supported by USDA National Institute of Food and Agriculture grant 2017-06481 (EVR, YPC and JDE) and the USDA Animal and Plant Health Inspection Service (APHIS). Genomic analyses used resources provided by the USDA-Agricultural Research Service SCINet project, ARS project number 0500-00093-001-00-D. USDA is an equal opportunity provider and employer. The authors declare that they have no competing interests.

## Author contribution

E.V.R. conceived and planned the study in consultation with J.D.E.; F.P., D.W. and D.vE. provided insect samples; E.V.R., D.L, K.G., W.G. carried out the laboratory molecular work. A.K.C. analyzed the NGS data; E.V.R, A.K.C., D.vE., Y.P.C, and J.D.E. wrote the manuscript. All co-authors contributed to data interpretation, and to the writing of the manuscript.

## Supporting information

**S1 Fig.**
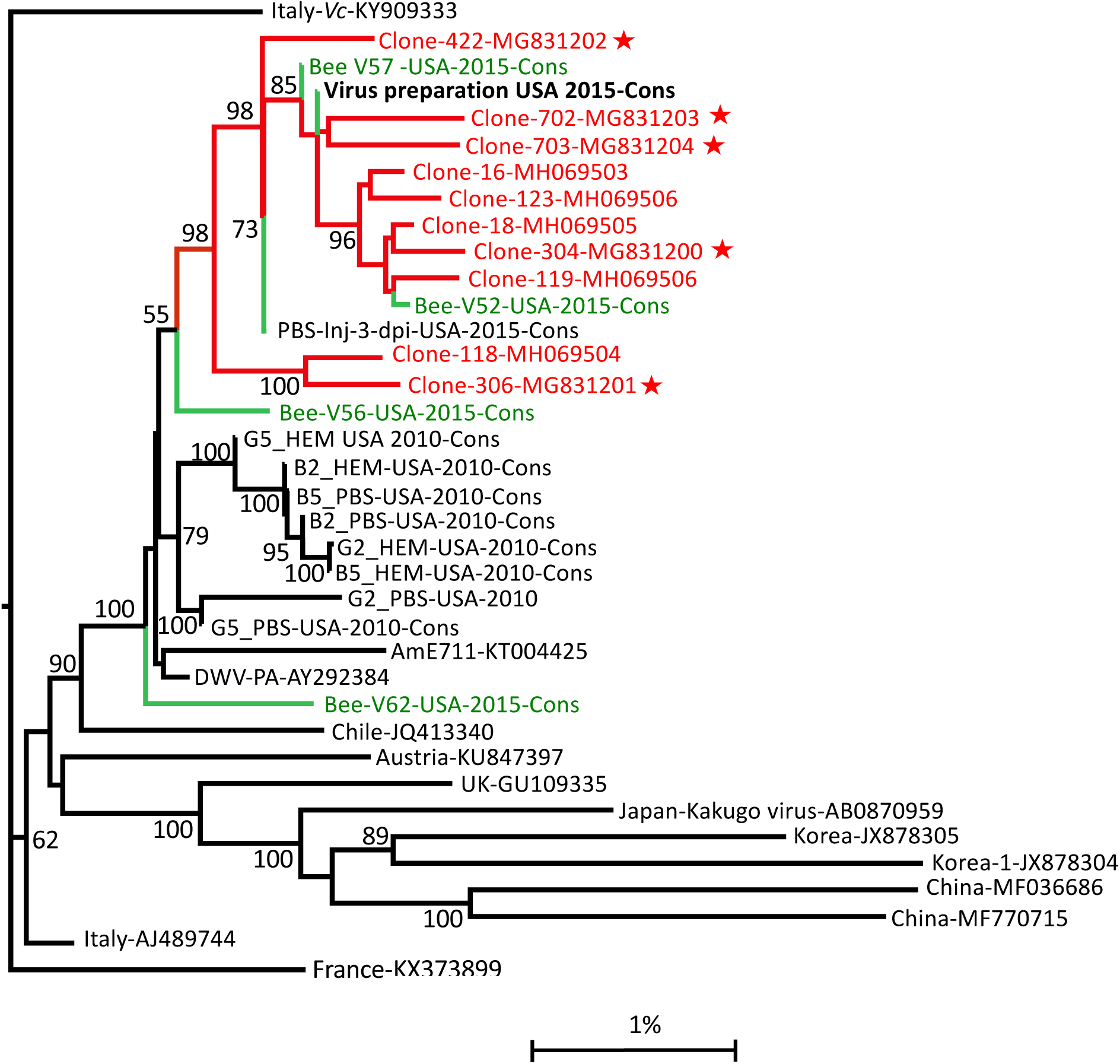
Phylogeny of the complete coding sequence of DWV genomic RNAs. A maximum likelihood phylogenetic tree was generated for a 9.7 kb DWV genome section (position 472 in the 3’ IRES to the 3’ poly(A) sequence, position 10162), which includes the entire protein-coding sequence. Bootstrap values for 1000 replicates are shown for the groups with more than 50% bootstrap support. Red nodes connect the cloned sequences from a Maryland *Varroa*-infested colony sampled in 2015 (“Clone-“). Stars indicate the DWV cDNA clones designed in this experiment and tested for infectivity. Suffix “-Cons” indicates consensus NGS sequences. Green nodes and labels indicate NGS consensus sequences for DWV from individual honey bee pupae. Scale shows genetic distance (%). Isolates are labelled with their clone identifier or country of origin as well as their NCBI accession.

**S2 Fig.**
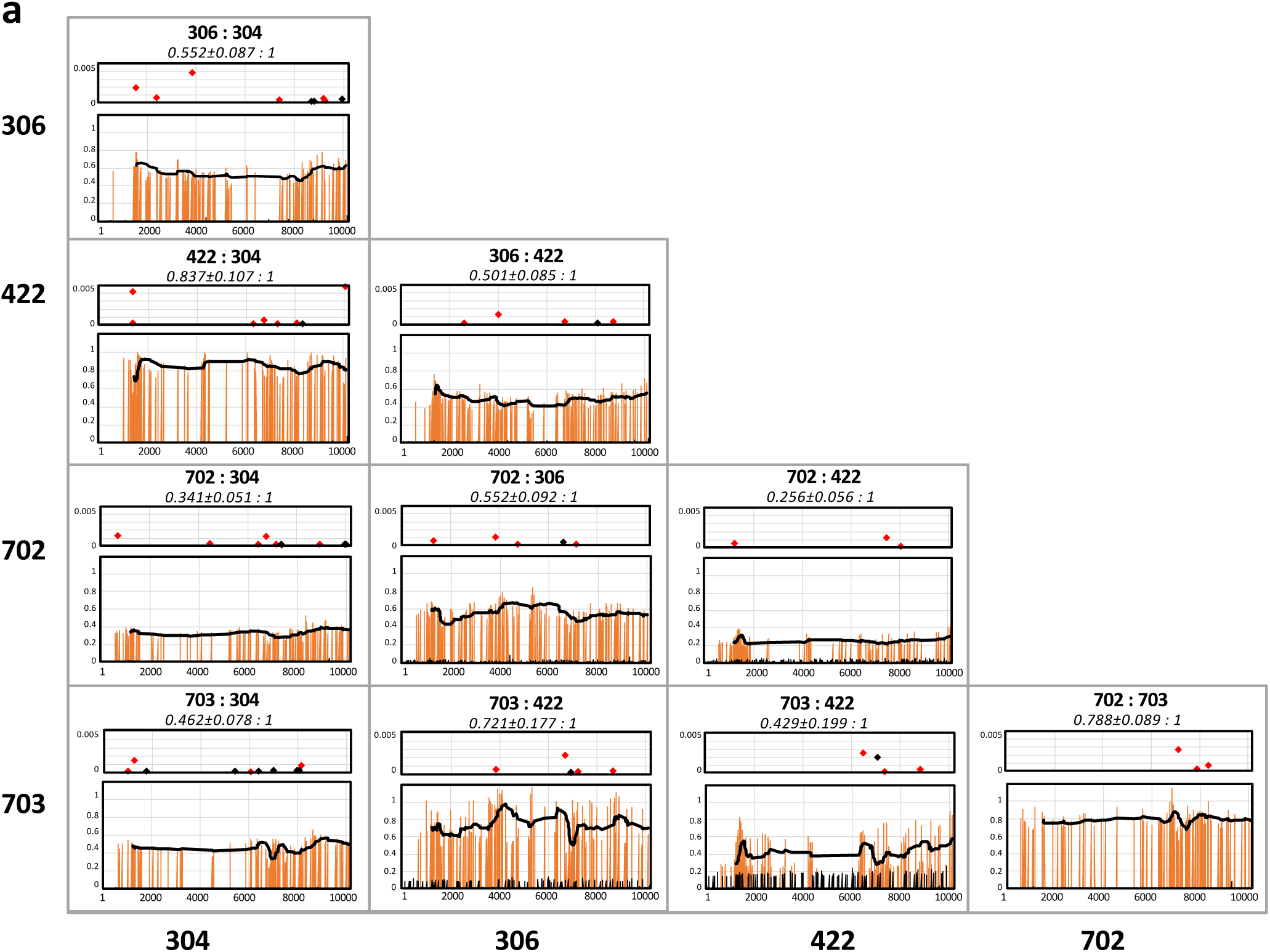
Analysis of the within-population competitiveness and recombination of DWV genotypes. Individual honey bee pupae were injected with two separate clone-derived DWV isolates, 5*10^6^ copies of each, and analyzed by NGS at 3 days post-injection (dpi). Overall ratios between the clone-derived DWV isolates in the progeny, calculated as an average of the expected polymorphic nucleotide proportions ± SD are shown for each combination. The top panels show recombination breakpoints identified using the LUMPY package, and their proportions in relation to the NGS read coverage. Only the recombination sites with more than 10 supporting events exceeding 0.02% of the NGS coverage are shown; black and red diamonds show the recombination sites (black - for the predominant isolate at 5’, red for the predominant isolate at 3’). In the lower panels the orange bars show ratios between the expected polymorphic nucleotides for each pair-wise combination along with trend lines. The black bars show the ratios of the second most abundant nucleotide to the total coverage at the positions, which were not polymorphic for a given genotype pair, indicative of the background wild-type DWV infection. For each graph, X-axes show nucleotide position in the DWV genome and Y-axes show recombinant read proportions (top panes) or polymorphic nucleotide ratios (bottom panes).

**S3 Fig.**
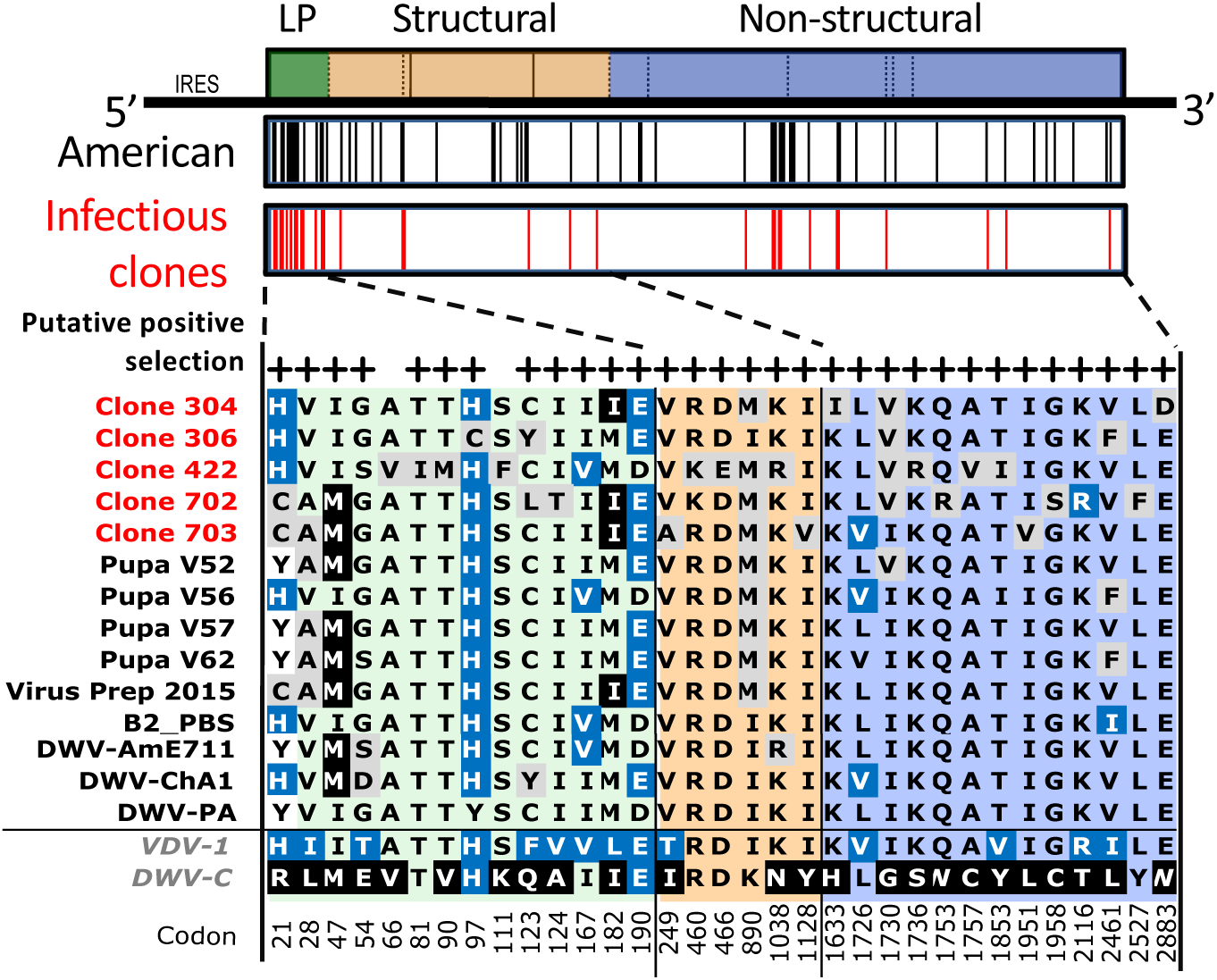
Diversifying selection of DWV proteins. Alignment shows divergent positions in the full-length American DWV sequences and in the infectious US DWV cDNA clones. Codon numbers are indicated below. Positions marked with “+” correspond to putative codons under positive diversifying selection in DWV strain from Pennsylvania (AY292384) is used as the reference genotype.

**S4 Fig.**
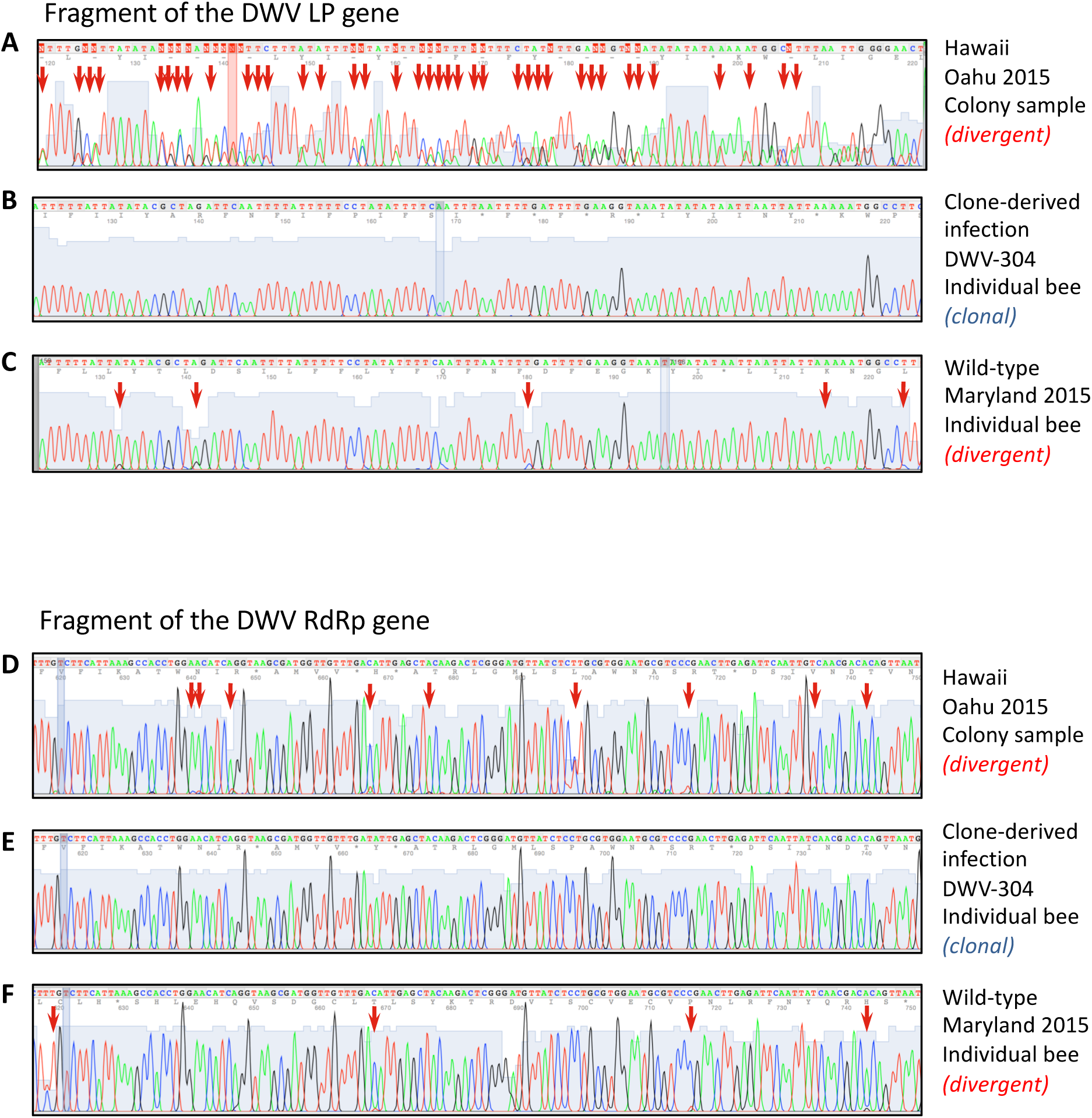
Polymorphisms in *Varroa-*associated DWV from 2015 Hawaii and Maryland honey bee samples. Electropherograms of the direct sequencing of RT-PCR fragments corresponding to sections of DWV RNA: (A-C) LP region (positions 1045-1158), and (D-F) RdRp fragment (positions 8645-8744; sequenced in the Hawaiian DWV diversity study to demonstrate DWV clonality in Oahu Island DWV in 2009 [9]). The fragments were amplified using RNA extracted from (A, D) pooled honey bees collected from a *Varroa-*infested apiary on Oahu Island, Hawaii, in 2015; (B, E) an individual honey bee pupa injected with clone-derived DWV-304; (C, F) an individual honey bee pupae infected with Maryland wild-type DWV (2015 sample source). Divergent positions are indicated with arrows.

**S5 Fig.**
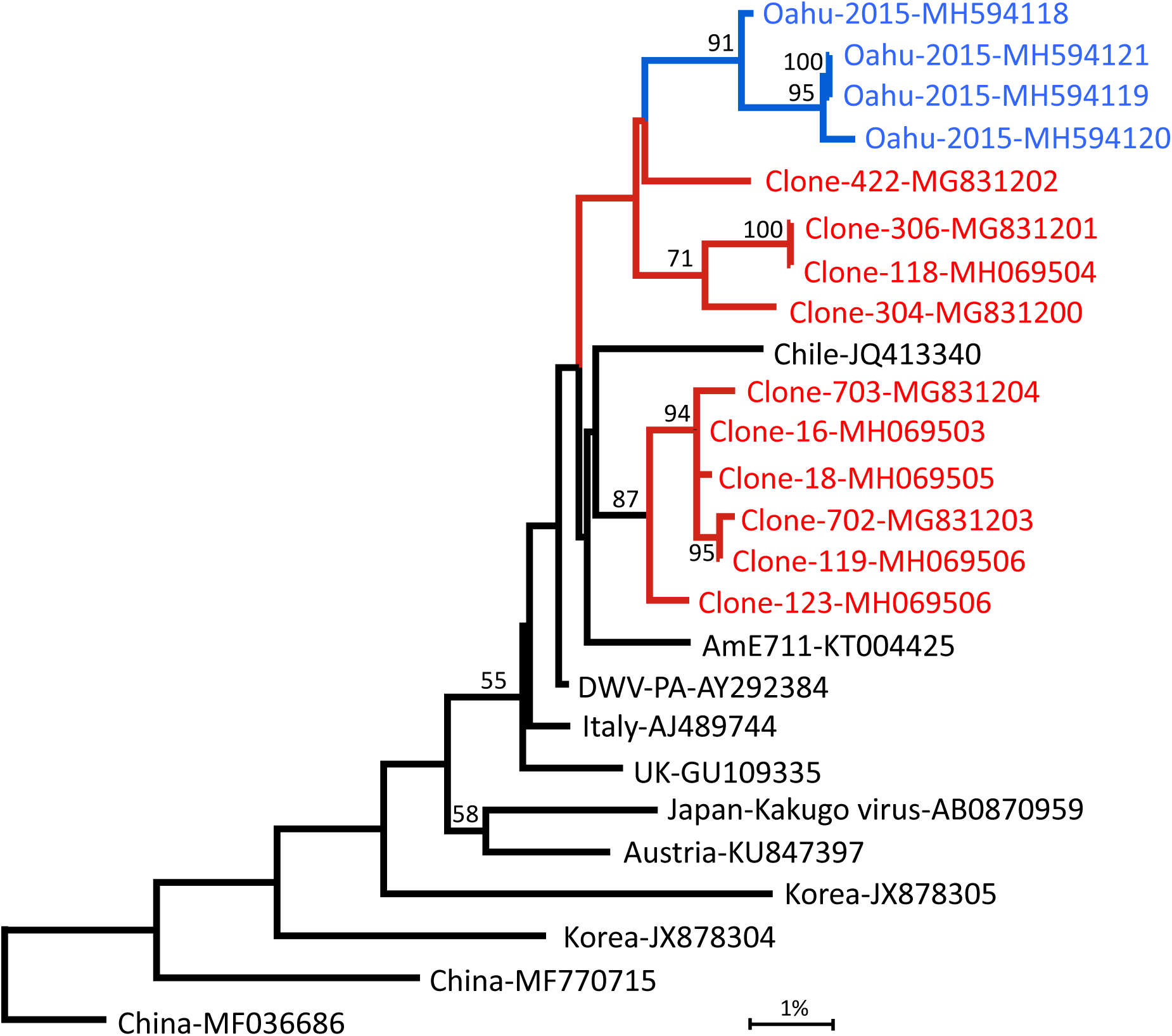
Phylogenetic analysis of the LP-coding sequences of DWV RNA. Maximum likelihood phylogenetic tree corresponds to the section of the DWV genome coding for the LP and adjacent parts of 5’ IRES and the structural gene block (positions 922 to 2048). Blue nodes connect the cloned sequences from a Hawaiian apiary (Oahu Island) sampled in 2015; red nodes connect the cloned sequences and the full-length DWV genomes from a Maryland *Varroa*-infested colony sampled in 2015. Bootstrap values for 1000 replicates are shown for the groups with more than 50% bootstrap support. Scale bar shows genetic distance (%). Isolates are labelled with their clone identifier or country of origin as well as their NCBI accession.

**S1 Text 1.**
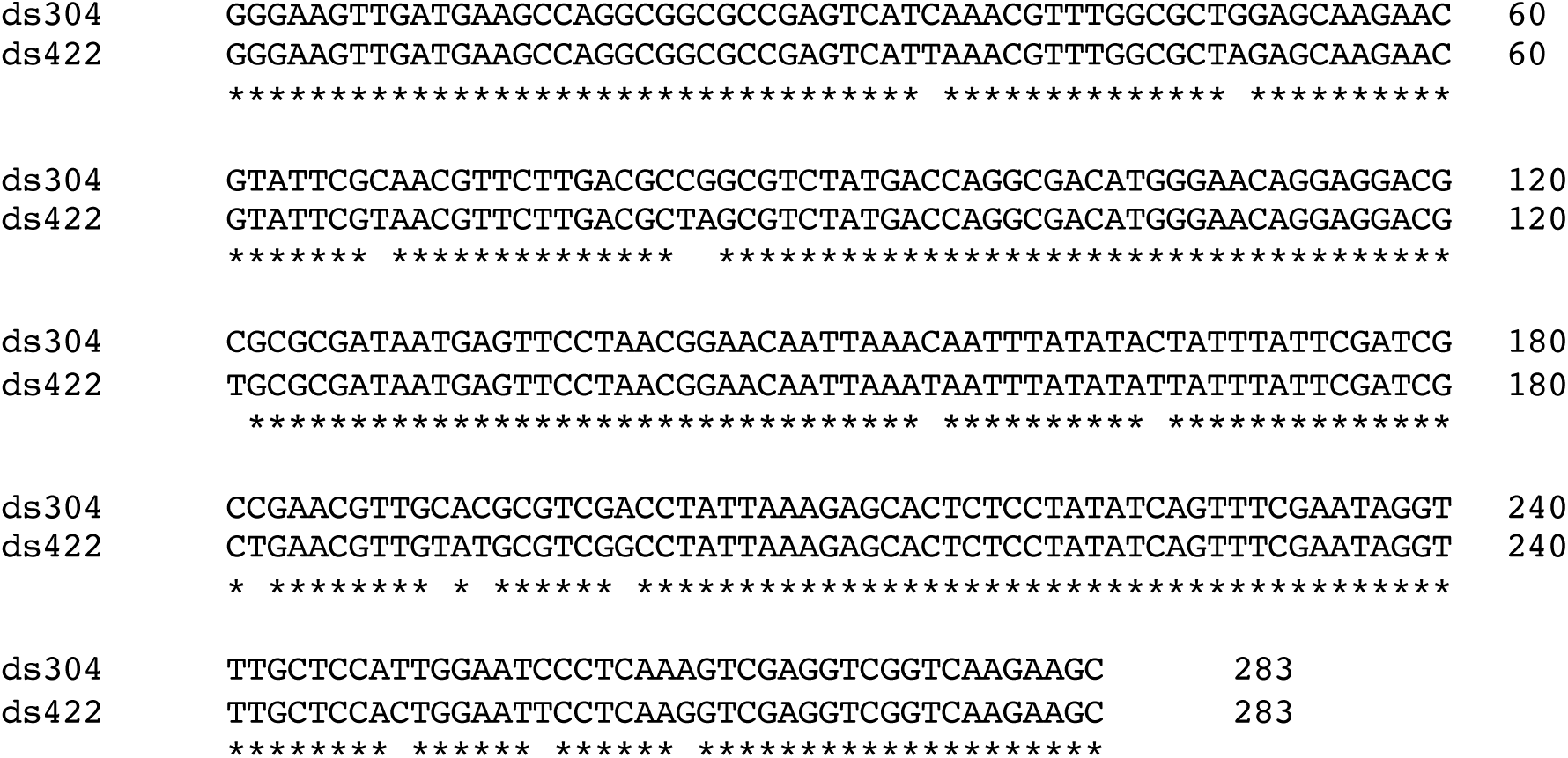
Double-stranded (ds) RNAs specific to the cloned DWV genotypes. Nucleotide alignment of the 283 nt section of DWV isolates DWV-304 and DWV-422, (Genbank accession numbers MG831200 and MG83120), positions 1242-1524 in the DWV genomic RNA, which were used to generate dsRNA.

**S1 Table.**
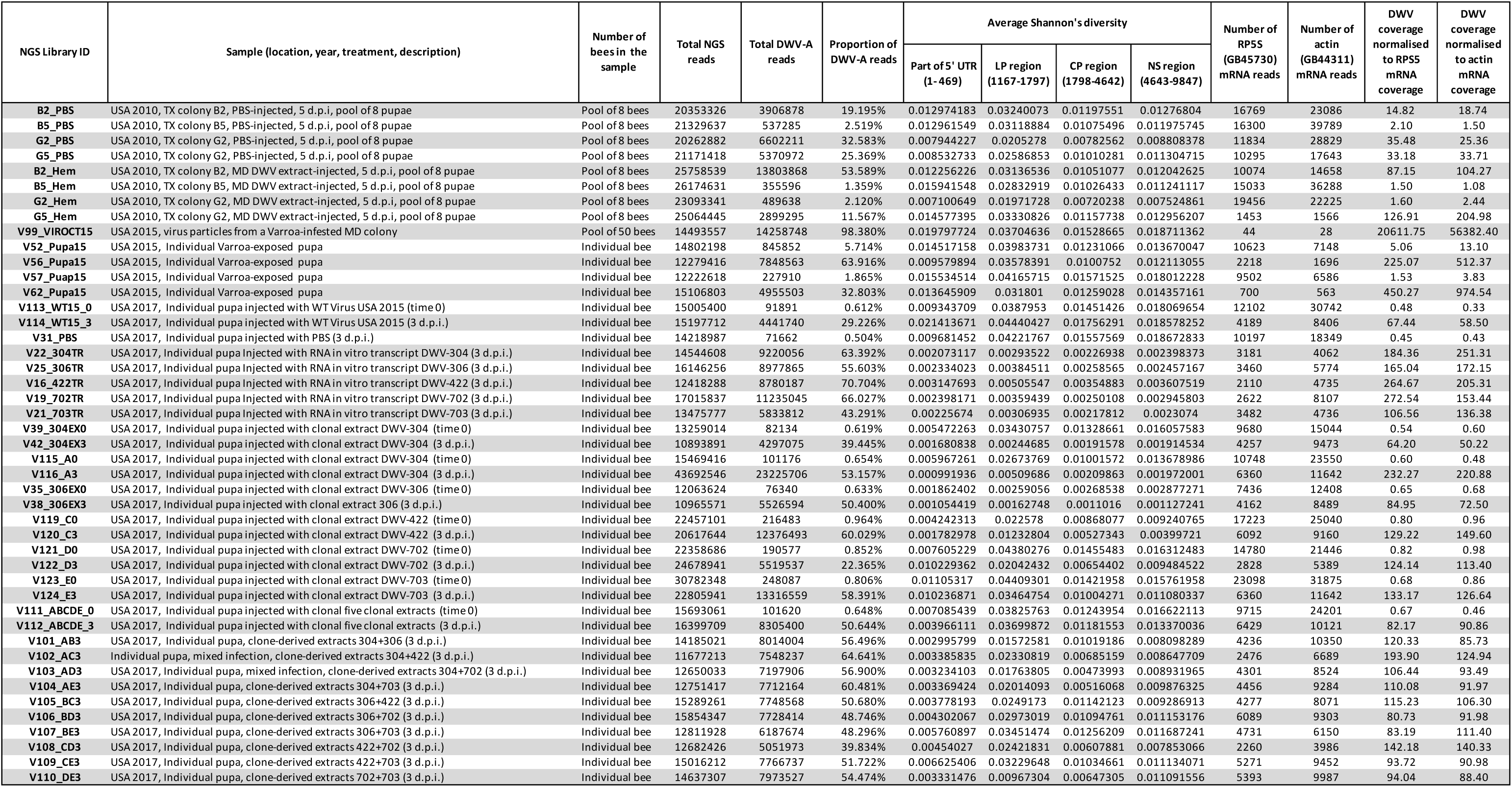
NGS libraries. NCBI Sequence Read Archive accession SRP135682, https://www.ncbi.nlm.nih.gov/sra/SRP135682.

**S2 Table.**
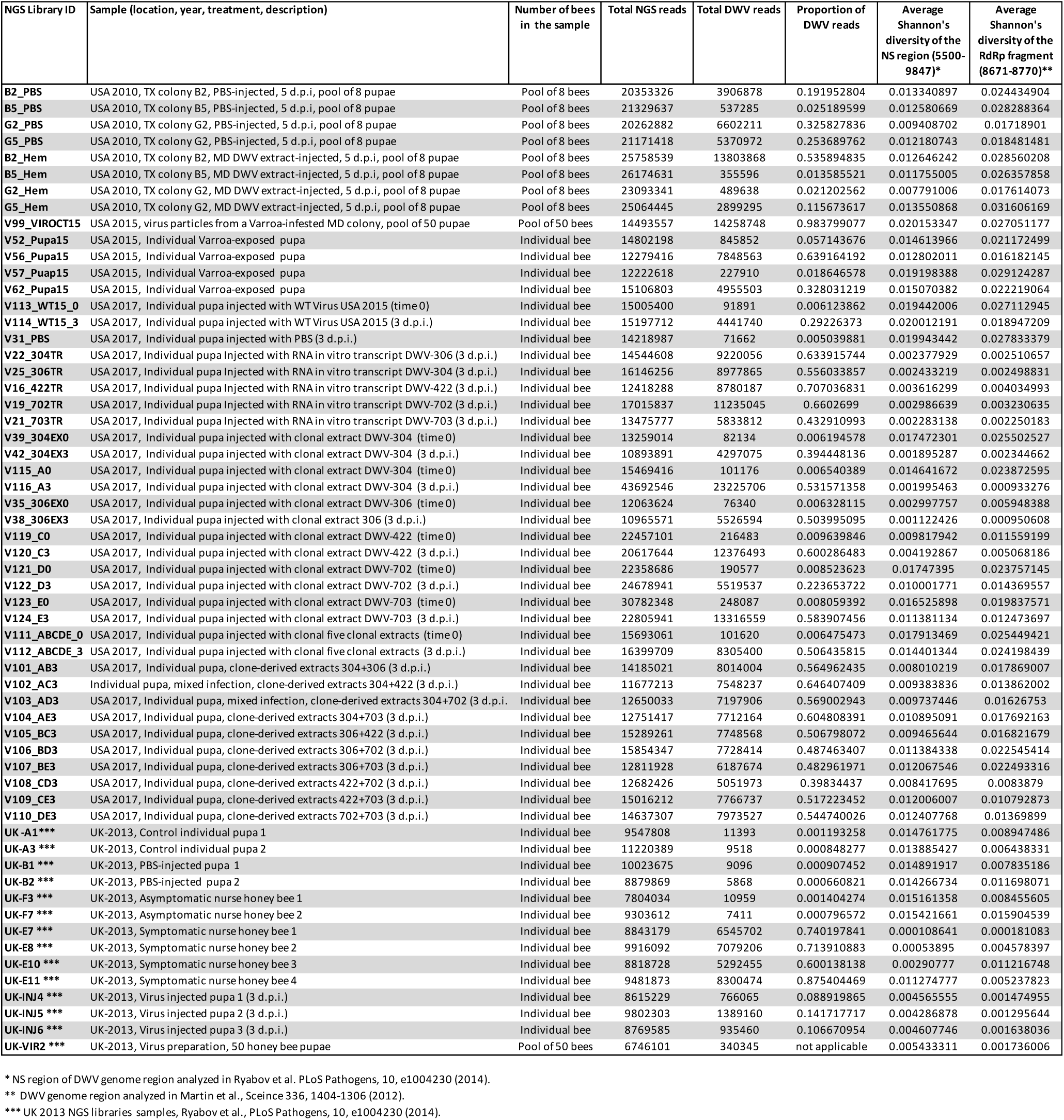
NGS summary and average Shannon’s diversity of the non-structural (NS) and RdRp regions of the US and UK DWV populations.

**S3 Table.** Comparison of the distribution of divergent nucleotides in DWV populations. Pearson correlation for Shannon’s diversity profiles for a 100 nt window. The degree of shade indicate the correlation coefficients: no shade - below 0.5, light shade - above 0.5 to 0.7, medium shade - from 0.7 to 0.9, dark shade - above 0.9.

| NGS library ID |  | B2_PBS | B5_PBS | G2_PBS | G5_PBS | B2_Hem | B5_Hem | G2_Hem | G5_Hem | V52_Pupa15 | V56_Pupa15 | V57_Puap15 | V62_Pupa15 | V99_VIROCT15 | V22_304TR | V25_306TR | V16_422TR | V19_702TR | V21_703TR | V39_304EX0 | V42_304EX3 | V31_PBS | V114_WT15_3 | V112_ABCDE_3 | Theoretic 5 cloned US DWV |
| --- | --- | --- | --- | --- | --- | --- | --- | --- | --- | --- | --- | --- | --- | --- | --- | --- | --- | --- | --- | --- | --- | --- | --- | --- | --- |
| B2_PBS | USA 2010, TX colony B2, PBS-injected, 5 dpi, pool of 8 pupae |  | 0.9684 | 0.9028 | 0.8256 | 0.9887 | 0.9623 | 0.9715 | 0.9840 | 0.7432 | 0.6998 | 0.7333 | 0.6639 | 0.6922 | 0.1144 | 0.2970 | 0.4020 | 0.1927 | 0.2485 | 0.5844 | 0.0842 | 0.6754 | 0.7090 | 0.6101 | 0.5813 |
| B5_PBS | USA 2010, TX colony B5, PBS-injected, 5 dpi, pool of 8 pupae | 0.9684 |  | 0.8970 | 0.8078 | 0.9770 | 0.9899 | 0.9474 | 0.9744 | 0.7331 | 0.6848 | 0.7285 | 0.6662 | 0.6828 | 0.1162 | 0.2929 | 0.4093 | 0.2047 | 0.2583 | 0.5880 | 0.1047 | 0.6645 | 0.6897 | 0.6024 | 0.5771 |
| G2_PBS | USA 2010, TX colony G2, PBS-injected, 5 dpi, pool of 8 pupae | 0.9028 | 0.8970 |  | 0.8938 | 0.9072 | 0.8840 | 0.8744 | 0.8968 | 0.6440 | 0.5857 | 0.6388 | 0.5867 | 0.6276 | 0.0847 | 0.2341 | 0.3484 | 0.1925 | 0.1939 | 0.5222 | 0.0540 | 0.5918 | 0.6289 | 0.5377 | 0.5188 |
| G5_PBS | USA 2010, TX colony G2, PBS-injected, 5 dpi, pool of 8 pupae | 0.8256 | 0.8078 | 0.8938 |  | 0.8227 | 0.8180 | 0.8137 | 0.8125 | 0.5941 | 0.5454 | 0.5878 | 0.5287 | 0.5666 | 0.0479 | 0.2254 | 0.3175 | 0.1920 | 0.1696 | 0.4647 | 0.0296 | 0.5407 | 0.6010 | 0.4976 | 0.4932 |
| B2_Hem | USA 2010, TX colony B2, MD DWV extract-injected, 5 dpi, pool of 8 pupae | 0.9887 | 0.9770 | 0.9072 | 0.8227 |  | 0.9726 | 0.9696 | 0.9962 | 0.7384 | 0.6925 | 0.7358 | 0.6694 | 0.6922 | 0.1041 | 0.2884 | 0.4034 | 0.1932 | 0.2523 | 0.5787 | 0.0818 | 0.6753 | 0.7031 | 0.6131 | 0.5869 |
| B5_Hem | USA 2010, TX colony B5, MD DWV extract-injected, 5 dpi, pool of 8 pupae | 0.9623 | 0.9899 | 0.8840 | 0.8180 | 0.9726 |  | 0.9647 | 0.9672 | 0.7027 | 0.6579 | 0.6992 | 0.6463 | 0.6452 | 0.1126 | 0.3046 | 0.4026 | 0.1988 | 0.2590 | 0.5580 | 0.1107 | 0.6360 | 0.6626 | 0.5790 | 0.5546 |
| G2_Hem | USA 2010, TX colony G2, MD DWV extract-injected, 5 dpi, pool of 8 pupae | 0.9715 | 0.9474 | 0.8744 | 0.8137 | 0.9696 | 0.9647 |  | 0.9592 | 0.6821 | 0.6544 | 0.6767 | 0.6309 | 0.6128 | 0.1136 | 0.3212 | 0.3992 | 0.1789 | 0.2485 | 0.5363 | 0.1170 | 0.6237 | 0.6394 | 0.5735 | 0.5452 |
| G5_Hem | USA 2010, TX colony G2, MD DWV extract-injected, 5 dpi, pool of 8 pupae | 0.9840 | 0.9744 | 0.8968 | 0.8125 | 0.9962 | 0.9672 | 0.9592 |  | 0.7484 | 0.7014 | 0.7413 | 0.6718 | 0.7017 | 0.1059 | 0.2884 | 0.4082 | 0.1931 | 0.2565 | 0.5751 | 0.0857 | 0.6765 | 0.7118 | 0.6122 | 0.5830 |
| V52_Pupa15 | USA 2015, Individual Varroa-exposed pupa | 0.7432 | 0.7331 | 0.6440 | 0.5941 | 0.7384 | 0.7027 | 0.6821 | 0.7484 |  | 0.8669 | 0.9549 | 0.8882 | 0.9158 | 0.2374 | 0.3713 | 0.4803 | 0.3428 | 0.3185 | 0.8098 | 0.1694 | 0.8643 | 0.8522 | 0.8034 | 0.7701 |
| V56_Pupa15 | USA 2015, Individual Varroa-exposed pupa | 0.6998 | 0.6848 | 0.5857 | 0.5454 | 0.6925 | 0.6579 | 0.6544 | 0.7014 | 0.8669 |  | 0.8293 | 0.7453 | 0.8053 | 0.2081 | 0.3039 | 0.4138 | 0.2662 | 0.2775 | 0.7263 | 0.1527 | 0.7775 | 0.7871 | 0.7186 | 0.6812 |
| V57_Puap15 | USA 2015, Individual Varroa-exposed pupa | 0.7333 | 0.7285 | 0.6388 | 0.5878 | 0.7358 | 0.6992 | 0.6767 | 0.7413 | 0.9549 | 0.8293 |  | 0.8868 | 0.9513 | 0.2602 | 0.3983 | 0.5136 | 0.4291 | 0.3700 | 0.8650 | 0.1836 | 0.9212 | 0.8480 | 0.8632 | 0.8378 |
| V62_Pupa15 | USA 2015, Individual Varroa-exposed pupa | 0.6639 | 0.6662 | 0.5867 | 0.5287 | 0.6694 | 0.6463 | 0.6309 | 0.6718 | 0.8882 | 0.7453 | 0.8868 |  | 0.8045 | 0.2708 | 0.3624 | 0.4809 | 0.3630 | 0.3349 | 0.7280 | 0.2597 | 0.7359 | 0.7113 | 0.6731 | 0.6391 |
| V99_VIROCT15 | USA 2015, virus particles from a Varroa-infested MD colony | 0.6922 | 0.6828 | 0.6276 | 0.5666 | 0.6922 | 0.6452 | 0.6128 | 0.7017 | 0.9158 | 0.8053 | 0.9513 | 0.8045 |  | 0.2425 | 0.3306 | 0.4436 | 0.4276 | 0.3329 | 0.8406 | 0.1254 | 0.8946 | 0.8820 | 0.8272 | 0.8041 |
| V22_304TR | USA 2017, Individual pupa injected with RNA in vitro transcript DWV-304 (3 dpi) | 0.1144 | 0.1162 | 0.0847 | 0.0479 | 0.1041 | 0.1126 | 0.1136 | 0.1059 | 0.2374 | 0.2081 | 0.2602 | 0.2708 | 0.2425 |  | 0.7846 | 0.7541 | 0.7664 | 0.8137 | 0.3360 | 0.8813 | 0.2703 | 0.1567 | 0.3012 | 0.2402 |
| V25_306TR | USA 2017, Individual pupa injected with RNA in vitro transcript DWV-306 (3 dpi) | 0.2970 | 0.2929 | 0.2341 | 0.2254 | 0.2884 | 0.3046 | 0.3212 | 0.2884 | 0.3713 | 0.3039 | 0.3983 | 0.3624 | 0.3306 | 0.7846 |  | 0.7376 | 0.7673 | 0.8376 | 0.4242 | 0.7374 | 0.4043 | 0.3005 | 0.4722 | 0.4406 |
| V16_422TR | USA 2017, Individual pupa injected with RNA in vitro transcript DWV-422 (3 dpi) | 0.4020 | 0.4093 | 0.3484 | 0.3175 | 0.4034 | 0.4026 | 0.3992 | 0.4082 | 0.4803 | 0.4138 | 0.5136 | 0.4809 | 0.4436 | 0.7541 | 0.7376 |  | 0.7062 | 0.7366 | 0.5220 | 0.6946 | 0.5251 | 0.3894 | 0.5552 | 0.5105 |
| V19_702TR | USA 2017, Individual pupa injected with RNA in vitro transcript DWV-702 (3 dpi) | 0.1927 | 0.2047 | 0.1925 | 0.1920 | 0.1932 | 0.1988 | 0.1789 | 0.1931 | 0.3428 | 0.2662 | 0.4291 | 0.3630 | 0.4276 | 0.7664 | 0.7673 | 0.7062 |  | 0.8197 | 0.4624 | 0.6970 | 0.4198 | 0.2965 | 0.4424 | 0.4286 |
| V21_703TR | USA 2017, Individual pupa injected with RNA in vitro transcript DWV-703 (3 dpi) | 0.2485 | 0.2583 | 0.1939 | 0.1696 | 0.2523 | 0.2590 | 0.2485 | 0.2565 | 0.3185 | 0.2775 | 0.3700 | 0.3349 | 0.3329 | 0.8137 | 0.8376 | 0.7366 | 0.8197 |  | 0.3744 | 0.7503 | 0.3635 | 0.2697 | 0.4118 | 0.3770 |
| V39_304EX0 | USA 2017, Individual pupa injected with clonal extract DWV-304 (time 0) | 0.5844 | 0.5880 | 0.5222 | 0.4647 | 0.5787 | 0.5580 | 0.5363 | 0.5751 | 0.8098 | 0.7263 | 0.8650 | 0.7280 | 0.8406 | 0.3360 | 0.4242 | 0.5220 | 0.4624 | 0.3744 |  | 0.2633 | 0.9239 | 0.7185 | 0.9004 | 0.8889 |
| V42_304EX3 | USA 2017, Individual pupa injected with clonal extract DWV-304 (3 dpi) | 0.0842 | 0.1047 | 0.0540 | 0.0296 | 0.0818 | 0.1107 | 0.1170 | 0.0857 | 0.1694 | 0.1527 | 0.1836 | 0.2597 | 0.1254 | 0.8813 | 0.7374 | 0.6946 | 0.6970 | 0.7503 | 0.2633 |  | 0.1602 | 0.0444 | 0.1986 | 0.1447 |
| V31_PBS | USA 2017, Individual pupa injected with PBS (3 dpi) | 0.6754 | 0.6645 | 0.5918 | 0.5407 | 0.6753 | 0.6360 | 0.6237 | 0.6765 | 0.8643 | 0.7775 | 0.9212 | 0.7359 | 0.8946 | 0.2703 | 0.4043 | 0.5251 | 0.4198 | 0.3635 | 0.9239 | 0.1602 |  | 0.8037 | 0.9378 | 0.9272 |
| V114_WT15_3 | USA 2017, Individual pupa injected with WT Virus USA 2015 (3 dpi) | 0.7090 | 0.6897 | 0.6289 | 0.6010 | 0.7031 | 0.6626 | 0.6394 | 0.7118 | 0.8522 | 0.7871 | 0.8480 | 0.7113 | 0.8820 | 0.1567 | 0.3005 | 0.3894 | 0.2965 | 0.2697 | 0.7185 | 0.0444 | 0.8037 |  | 0.7372 | 0.6949 |
| V112_ABCDE_3 | USA 2017, Individual pupa injected with clonal five clonal extracts (3 dpi) | 0.6101 | 0.6024 | 0.5377 | 0.4976 | 0.6131 | 0.5790 | 0.5735 | 0.6122 | 0.8034 | 0.7186 | 0.8632 | 0.6731 | 0.8272 | 0.3012 | 0.4722 | 0.5552 | 0.4424 | 0.4118 | 0.9004 | 0.1986 | 0.9378 | 0.7372 |  | 0.9825 |
| Theoretical 5 cloned US DWV | Theoretical profile for the five cloned US DWV isolates, equal proportions of each | 0.5813 | 0.5771 | 0.5188 | 0.4932 | 0.5869 | 0.5546 | 0.5452 | 0.5830 | 0.7701 | 0.6812 | 0.8378 | 0.6391 | 0.8041 | 0.2402 | 0.4406 | 0.5105 | 0.4286 | 0.3770 | 0.8889 | 0.1447 | 0.9272 | 0.6949 | 0.9825 |  |

**S4 Table.**
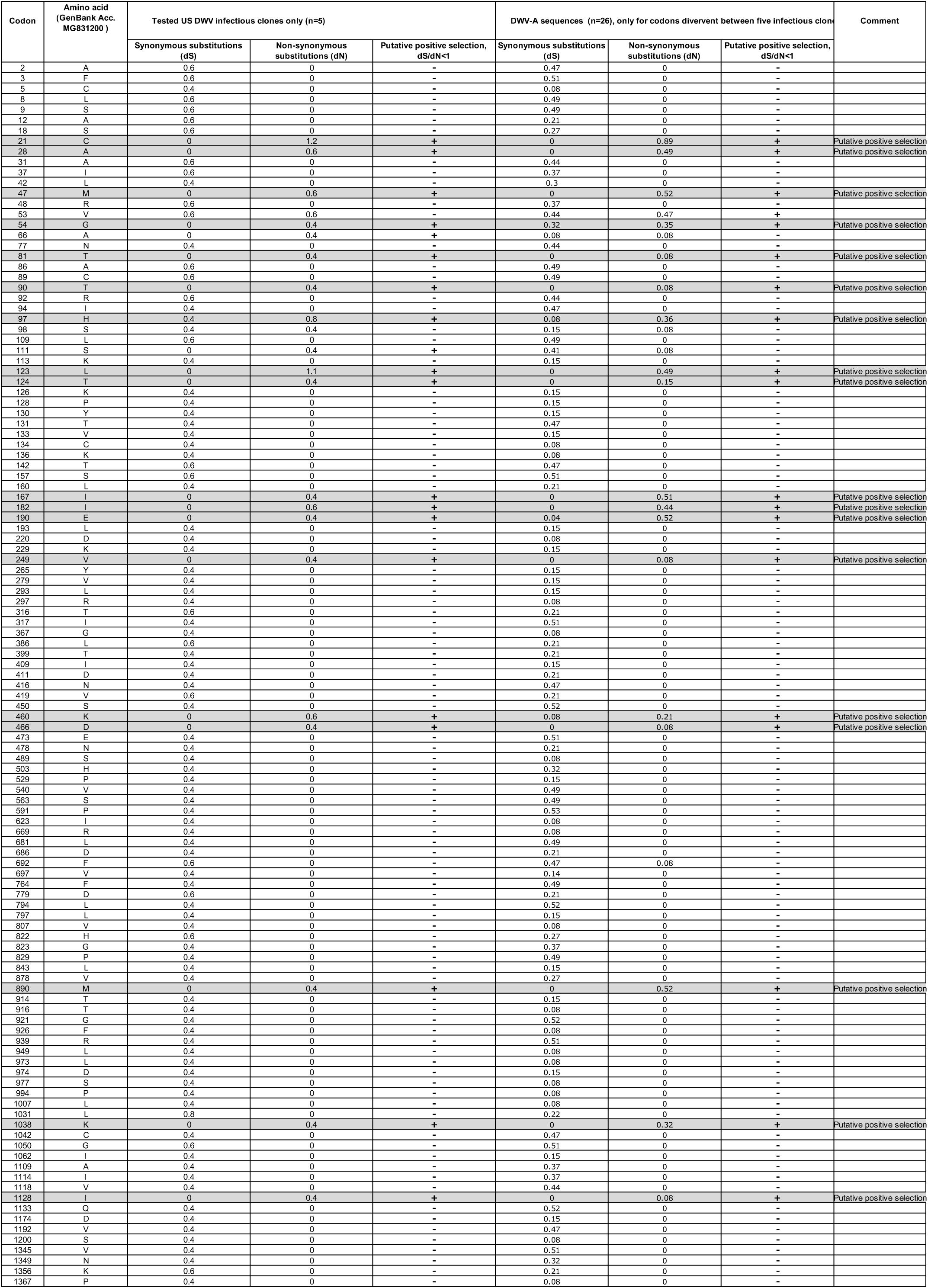
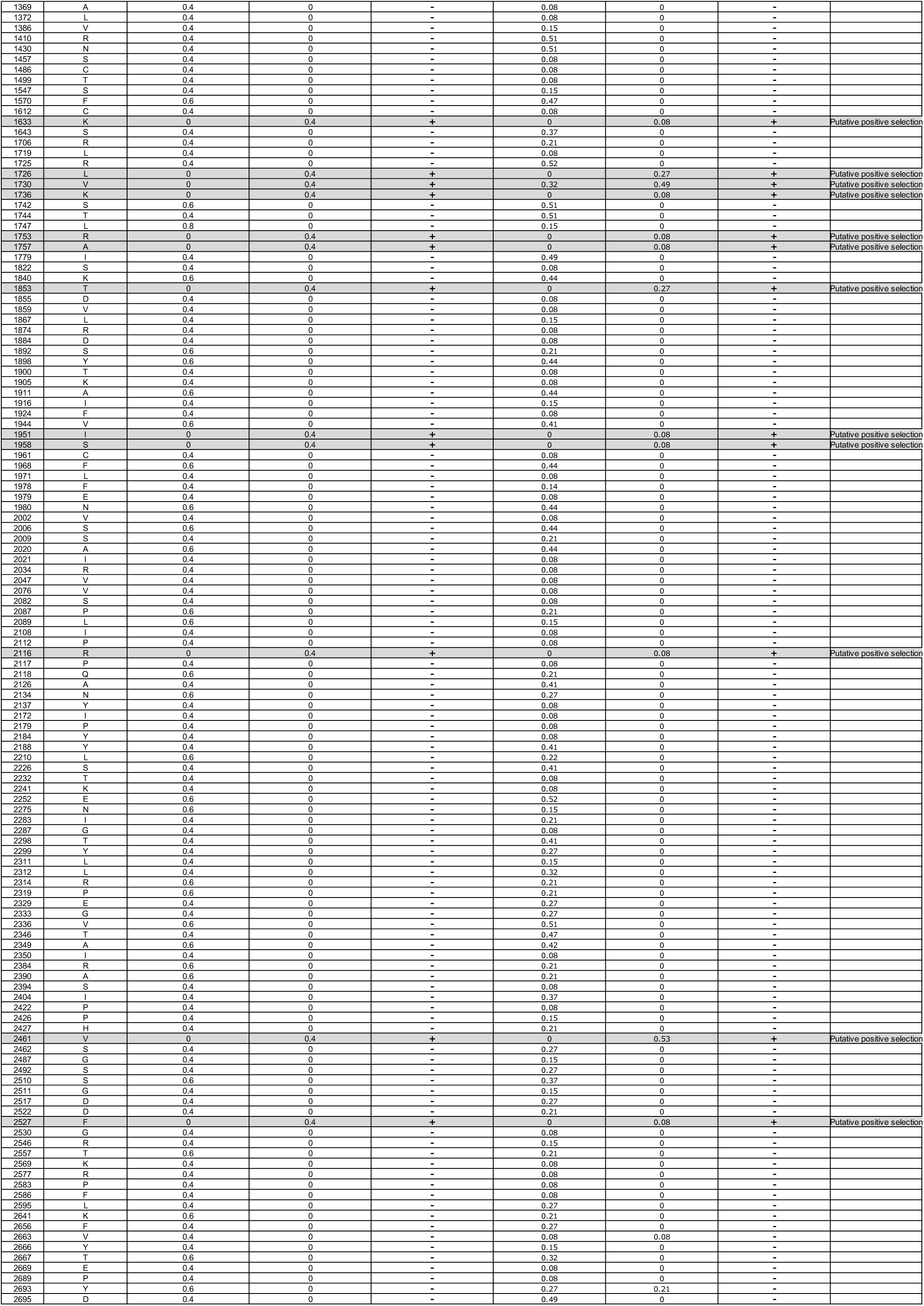
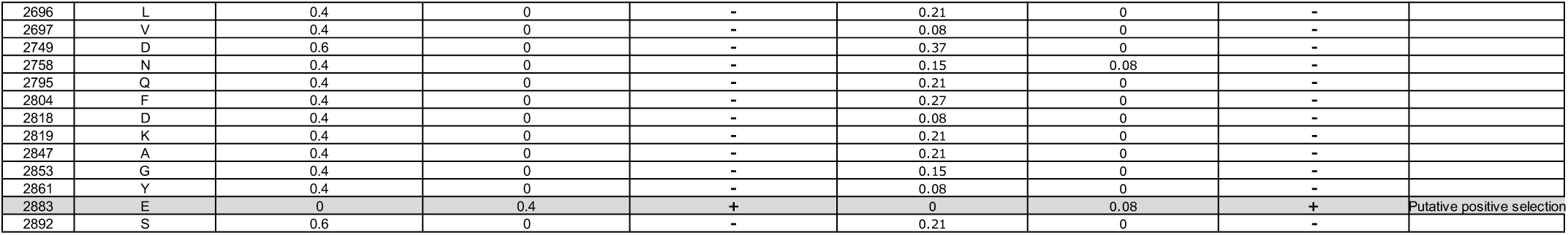
Synonymous and nonsynonymous substitutions in the infectious DWV clones.

**S5 Table.**
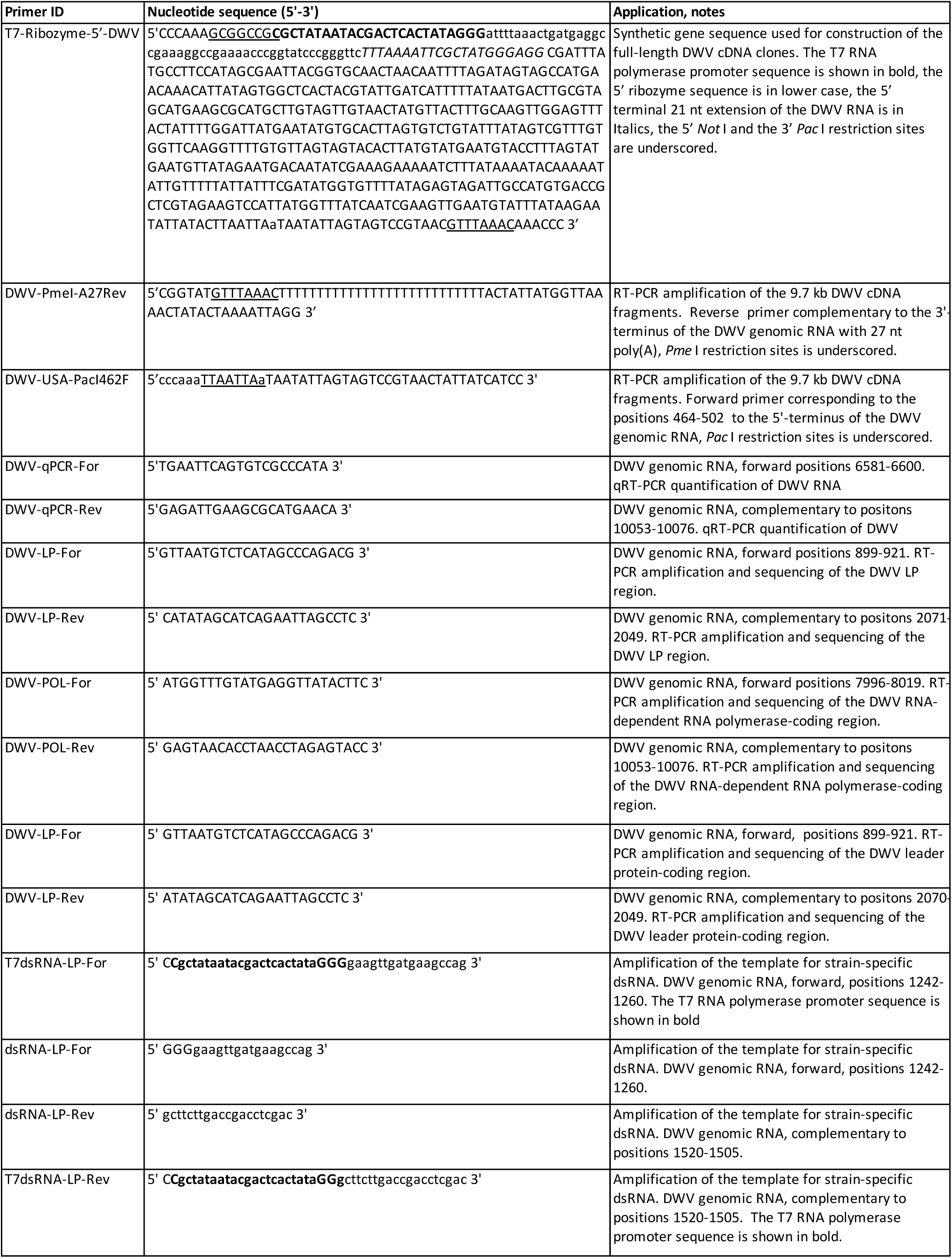
Primers and the synthetic gene used in this study.

